# From theory to application: Elasticity-consistent aggregation of Leslie matrix population models for comparative demography

**DOI:** 10.64898/2026.02.04.703802

**Authors:** Richard A. Hinrichsen, Hiroyuki Yokomizo, Roberto Salguero-Gómez

## Abstract

1. Ecology has entered the big data era. This influx of data has now enabled ecologists to address questions about life history and demographic patterns across the tree of life. Supporting this momentum, the COMADRE and COMPADRE databases represent a boon to comparative demography. However, these initiatives also present challenges due to the complexities of the life cycles that they describe. Matrix population models can vary in sampling frequency, life cycle stage width, life cycle complexity, and census type (e.g. pre- or post-reproduction). Complicating this picture is the fact that there are myriad matrix model representations of the same population, and model representation influences key demographic parameters. Thus, a key challenge in exploiting large demographic datasets is fair comparison of models constructed with different projection intervals and complexities.
2. One way to compare models of different complexity is to reduce (‘aggregate’) the larger models to the dimensionality of the smaller models. The commonly used aggregator (i.e. the standard aggregator), although it yields stable growth rate and stable stage distribution consistent with the original demographic model, does not provide consistent reproductive values and elasticities. To address this limitation, we extend and validate an existing elasticity-consistent aggregator, overcoming several of its methodological and biological limitations. Specifically, we derive the aggregator using balancing and interstage flows, which removes the requirement of matrix primitivity, and we restrict aggregation to Leslie-to-Leslie models, thereby preventing biologically infeasible survival probabilities exceeding one. We further introduce explicit metrics of aggregation effectiveness and apply the approach to 12 Leslie matrix population models representing animal populations from diverse taxonomic classes.
3. The elasticity-consistent aggregator returns a Leslie matrix that preserves key properties of the original matrix, including irreducibility and primitivity, and yields consistent estimates of population growth rate, stable age distribution, and reproductive values. Moreover, across all aggregated models from the 12 examined animal populations, the elasticity-consistent aggregator produced more accurate estimates than the standard aggregator for 86% of generation times, 60% of Demetrius’ entropies, and 76% of net reproductive rates.
4. By preserving key properties of the original model, the elasticity-consistent aggregator provides a useful framework for comparing matrix population models of varying complexity in comparative demography. Such a method helps seize the promise of big data in ecology and discover principles across the tree of life, from microbes to fungi, plants, and animals.

## 1. INTRODUCTION

The advent of big data in ecology allows us to address questions at ever larger spatial scales and over longer time frames (Hampton et al., 2013). For population ecology, this new era of big data means that we can now address a variety of questions about life history and demographic patterns across a wide variety of species (Salguero-Gómez & Gamelon, 2021). A stark example is the identification of the slow-fast continuum as the dominant axis of life history variation, a pattern revealed with striking clarity by large demographic datasets and supported across mammals (Gaillard et al., 1989; Oli & Dobson, 2003; Stearns, 1983), birds (Gaillard et al., 1989), insects (Blackburn, 1991; Bakewell et al., 2020), fish (Rochet, 2000), and plants (Salguero-Gómez et al., 2016b). Other studies using big data have revealed important patterns, such as the diversity of ageing across the tree of life (Jones et al., 2014), and the role of generation time in modulating climate responses of herbaceous perennial plants (Compagnoni, 2021).

In the last decade, comparative demography has surged as a main area of population ecology thanks to open-access EltonTraits (Wilman et al., 2014) for foraging attributes, and PADRINO integral projection model database (Levin et al., 2022) and the COMADRE and COMPADRE matrix population model (MPM) databases (Salguero-Gómez et al., 2015, 2016a) for demography. However, these efforts have drawbacks because the data they contain are author-driven (i.e. shaped by the individual choices and assumptions of the original authors) and heterogeneous in complexity (Che-Castaldo et al., 2020; Gascoigne et al., 2023; Kendall, 2015; Römer et al., 2024; Salguero-Gómez et al., 2021). Figure 1 illustrates the diversity of matrix population models in the COMADRE database with respect to dimensionality and projection interval (time step). Leslie matrices (Leslie, 1945), where individuals in a population are structured by age, vary in the resolution of their schedules of vital rates (i.e. survival and reproductive rates) according to the chosen time step (Caswell, 2001). Leslie matrices can also vary by census time: immediately before (pre-reproductive census) or immediately after (post-reproduction census) reproduction (Caswell, 2001; Vindenes et al., 2021). Different choices for time step, age class definitions, and census time can alter the Leslie matrix, resulting in many possible variations of the matrix for the same population or species. For example, Leslie matrices for the Asian elephant (*Elephas maximus*) were constructed with differing age class widths of 1 year and 5 years, which leads to matrix dimensionalities of 60 and 12, respectively (Chelliah et al., 2013; Goswami et al., 2014). As such, comparing the highly variable Leslie matrix models that exist in the peer-reviewed literature can be challenging. In fact, stage/age width influences estimates of population growth rate, elasticities, life history traits, etc. (e.g. Enright et al., 1995; Picard & Liang, 2014; Ramula & Lehtilä, 2005; Salguero-Gómez & Plotkin, 2010). Consequently, variation in MPM construction represents an important source of bias in comparative analyses.

**FIGURE 1.**
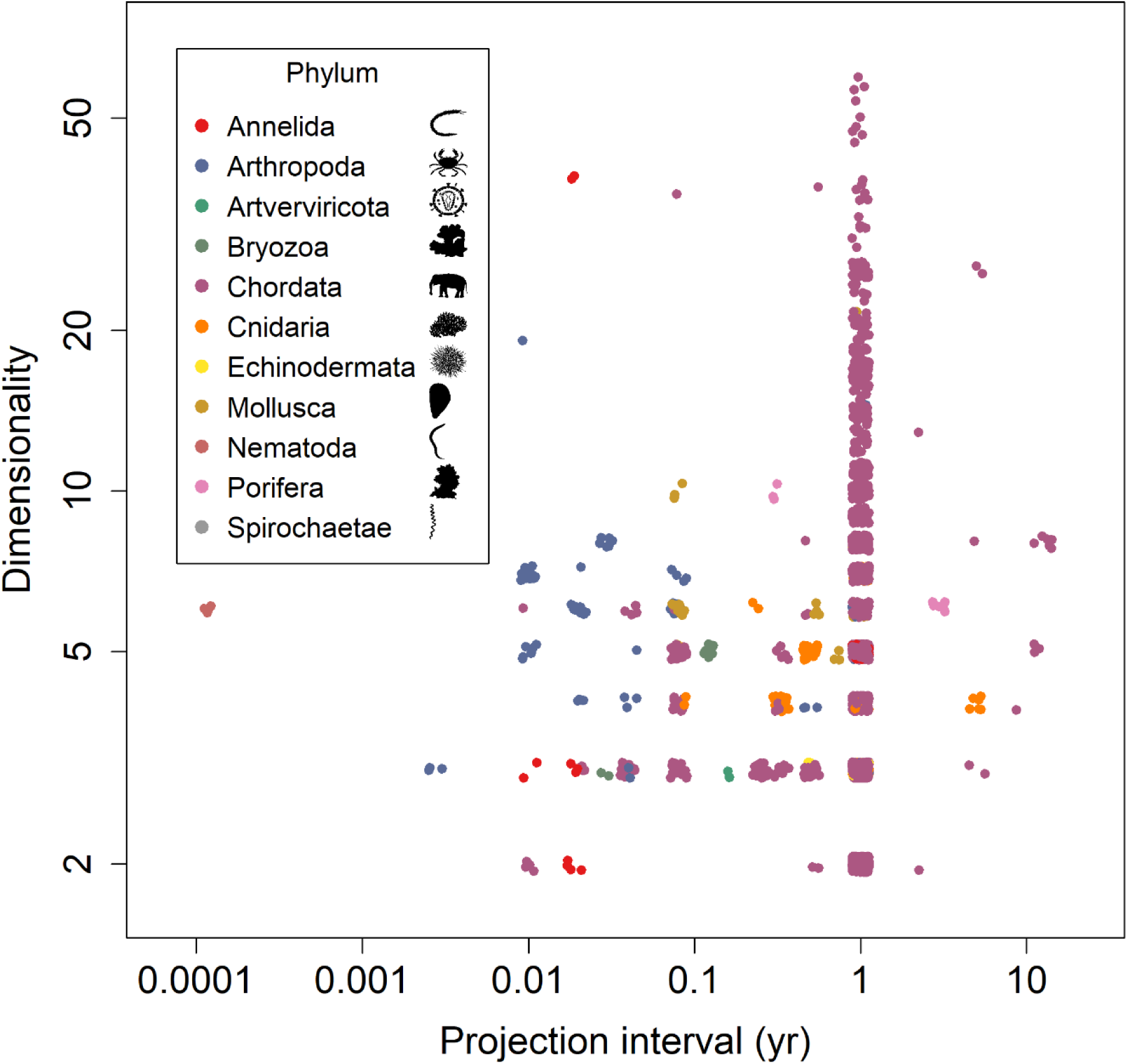
The rich variety of matrix population models (MPMs) in the COMADRE Animal Matrix Database (version 4.23.3.1) illustrated by the projection interval (the time step of each matrix population model) and the dimensionality (the number of stages in the model) across 11 animal phyla. Both axes are plotted on a log_10_ scale, and jitter was applied to separate overlapping observations. The heterogeneity of the MPMs makes comparative demography challenging. The silhouettes are from PhyloPic (https://www.phylopic.org; Keesey, 2023) and were added using the rphylopic R package (Gearty & Jones, 2023). Silhouettes were contributed as follows: Eunicidae by Thais Kananda da Silva Souza (2023; PDM 1.0); *Bythograea* by Kurtis Wothe & Giovanna Sainz (2024; CC0 1.0); HIV-1 by Arcadia Science (2025; CC0 1.0); *Celleporina grandis* by Guillaume Dera (2023; CC0 1.0*); Elephas maximus* by Andy Wilson (2023; CC0 1.0); *Acropora hyacinthus* by Guillaume Dera (2023; CC0 1.0); *Strongylocentrotus purpuratus* by Christoph Schomburg & Kurtis Wothe (2024; CC0 1.0); *Mytilus galloprovincialis* by Alexis Simon (2020; CC0 1.0); *Caenorhabditis elegans* by Taylor Medwig-Kinney (2023; CC0 1.0); *Spongia officinalis* by Guillaume Dera (2023; CC0 1.0); and *Arthrospira platensis* by Guillaume Dera (2022; CC0 1.0).

To compare two MPMs of different dimensionalities, one might aggregate the larger MPM so that its classes coincide with those of the smaller MPM, making vital rates and demographic parameters (e.g. generation time, life expectancy) directly comparable (Enright et al., 1995; Salguero-Gómez & Plotkin, 2010). Aggregation techniques, whereby one reduces a model to one of lower dimensionality, are well-developed for input-output analysis in economics (Howe & Johnson, 1989; Ijiri, 1971), automatic control (Sinha & Kuszta, 1983), and ecology (Auger et al., 2008; Iwasa et al., 1987; Iwasa et al., 1989). Several authors have developed the theory to aggregate multiregional MPMs into single-region MPMs (Auger et al., 2008; Sánchez et al., 1995; Sanz & Bravo de la Parra, 1999). Using the ‘standard aggregator’ (hereafter), others have aggregated age or stage classes of MPMs to form new classes that are consequently wider (Hinrichsen, 2023; Hooley, 2000; Salguero-Gómez & Plotkin, 2010).

A difficulty with the standard aggregator is that, although it produces stable (long-term) growth rate and stable age (or stage) distributions that are consistent with the original model (Hinrichsen, 2023; Hooley, 2000; Salguero-Gómez & Plotkin, 2010), it yields inconsistent reproductive values (i.e. the expected individual contributions to future generations *sensu* Fisher (1930)) (Bienvenu et al., 2017). This inconsistency, in turn, leads to inconsistent elasticities (i.e. relative contributions of vital rates to stable growth rate) (Bienvenu et al., 2017; Salguero-Gómez & Plotkin, 2010), which play a vital role in comparative demography (Franco & Silvertown, 2004), conservation biology (Menges, 2000), and evolutionary biology (van Tienderen, 1995, 2000).

Estimation of other demographic parameters can also rely on reproductive values, such as generation time, that is the average age at which individuals reproduce (Bienvenu & Legendre, 2015); inertia, a measure of a population’s eventual size relative to that expected if it had remained at its stable structure after a disturbance (Koons et al., 2007); population momentum, a variant of inertia (Caswell, 2001); and the sensitivity of the stable population growth rate to changes in vital rates (Caswell, 2001; de Kroon et al., 1986). Furthermore, reproductive value is the fundamental quantity maximized in optimizations of life histories (Charlesworth, 1994; Fisher, 1930; Goodman, 1982). The foundational role of reproductive values suggests that an aggregator that produces consistent reproductive values in addition to stable growth rate and stable age distribution offers a distinct advantage.

Here, we extend the ‘genealogical collapse’ method of Bienvenu et al. (2017), which we refer to as an ‘elasticity-consistent’ (hereafter) aggregator, to improve its practical applicability for comparative demography using matrix population models (MPMs). In its original form, it yields consistent reproductive values and elasticities together with consistent population growth rates and stable age distributions. However, the framework of Bienvenu et al. (2017) has three limitations that hinder its practical use in comparative demography: it is restricted to MPMs with primitive projection matrices (matrices with a strictly dominant positive eigenvalue), it can produce survival probabilities that exceed one, and quantitative diagnostics of aggregation error are absent. To relax the assumption of matrix primitivity, we re-derive the aggregator using balancing and a new ‘method of interstage flows’ (hereafter), which provides a shortcut to weighted least squares estimation (Hinrichsen 2023; Iwasa et al. 1989). Interstage flows quantify the movement of individuals between stages (Yokomizo et al. 2024) and enable aggregation of imprimitive matrices, such as those describing the bog fritillary butterfly (*Proclossiana eunomia*). To address the issue of survival probabilities that exceed one, we restrict aggregation to Leslie MPM → Leslie MPM. Finally, to address the lack of aggregation diagnostics, we introduce explicit aggregation-effectiveness measures that quantify aggregation error (Hinrichsen 2023; Iwasa et al. 1989).

We assess the elasticity-consistent aggregator’s performance by comparing its accuracy with that of the standard aggregator for three core demographic parameters: generation time; Demetrius’ entropy, which measures the spread of reproduction across ages (Caswell, 2001; Demetrius, 1974); and net reproductive rate, the average number of offspring produced by an individual over its expected lifetime (Caswell, 2001). The evaluation uses 12 age-based MPMs drawn from distinct taxonomic classes in the COMADRE Animal Matrix Database (Salguero-Gómez et al., 2016a), thus testing for the potential broad taxonomic applicability of our aggregator.

## 2. METHODS

### 2.1 Leslie MPM aggregation

A Leslie MPM is governed by the *n* × *n* Leslie matrix of the form

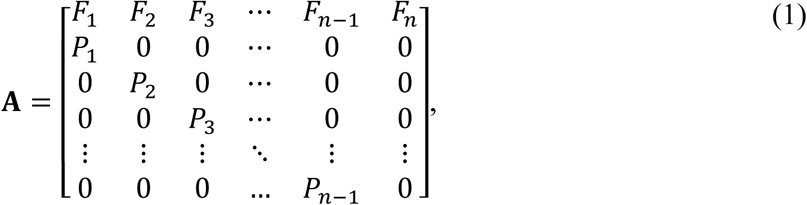

where *n* is the number of age classes, *F*_1_, *F*_2_, … , *F*_*n*_ are the age-specific fertility rates and *P*_1_, *P*_2_, … , *P*_*n*−1_ are the age-specific survival probabilities, which are assumed to be positive (Caswell, 2001; Leslie, 1945). We assume that **A** is irreducible (i.e. there is a path from any age class to every other), which, by the Perron-Frobenius Theorem, guarantees that stable age distribution, **w**, and the reproductive values, **v** (scaled so that **v**^⊤^**w** = 1), are both positive (Caswell, 2001; Freese & Johnson, 1974). A Leslie matrix is irreducible precisely when the fertility rate of the oldest age class is positive (Freese & Johnson, 1974).

To aggregate, we represent the original Leslie MPM governed by *n* × *n* Leslie matrix **A** with a smaller aggregated Leslie MPM governed by *m* × *m* Leslie matrix **B**, which is known as the aggregated matrix (Hinrichsen, 2023). Here we assume that *n* is a multiple of *m* (i.e. *n* = *km*, where *k* is a natural number), an assumption we can later relax. We partition the age classes of **A** into *m* wider age classes, each consisting of *k* adjacent age classes that form the age classes of the aggregated matrix **B**. The aggregated **B** satisfies

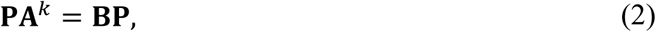

where the *m* × *n* matrix **P**, known as a partitioning matrix (*sensu* Howe & Johnson, 1989), is

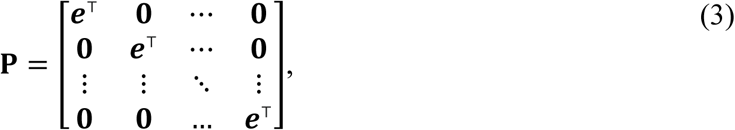

with **e** a *k*-vector of ones. The matrix **P** sums adjacent age classes, and **A**^*k*^ appears because **B** projects the population forward by *k*Δ*t*, where Δ*t* is the projection interval of the original MPM.

If **B** satisfies Equation (2) exactly, there is perfect aggregation of **A**^*k*^ associated with **P** and exact consistency with the original model (Howe & Johnson, 1989; Iwasa et al., 1987). Because Equation (2) is overdetermined (more equations than unknowns), an exact solution generally does not exist, and **B** is obtained by weighted least squares (Hinrichsen, 2023).

### 2.2 Standard aggregator

To obtain the standard aggregator, we minimize the weighted sum of squared errors,

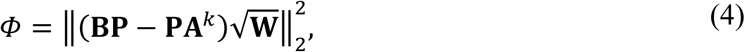

where ‖·‖_2_ is the Frobenius matrix norm, equal to the square root of the sum of squared entries (Horn & Johnson, 2013), and **W** = diag(**w**) is the *n* × *n* weight matrix, where diag(·) is the square diagonal matrix with diagonal equal to the indicated vector (in this case, the stable age distribution). Following Hinrichsen (2023), the solution is

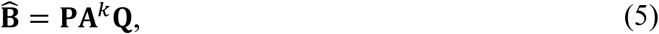

where **Q** is the *n* × *m* matrix

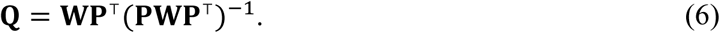

This method agrees with that of Hooley (2000) who aggregated **A** instead of **A**^*k*^ and used a more general partitioning matrix. But our choice to aggregate **A**^*k*^ with partitioning matrix **P**, guarantees that **B̂** is a Leslie matrix (Hinrichsen, 2023).

#### 2.2.1 Method of interstage flows

The method of interstage flows provides a shortcut to weighted least squares. The interstage flow of a population projection matrix **M**, denoted ℱ(**M**), is defined as **M** multiplied on the right by a diagonal matrix whose diagonal is the stable age distribution (Yokomizo et al., 2024). Aggregation equates interstage flows of the aggregated matrix with summed interstage flows of **A**^*k*^,

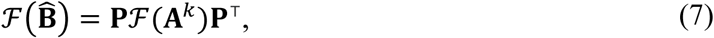

which can be written as

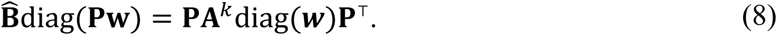

Solving this equation yields **B̂** = **PA**^*k*^**Q**, identical to the weighted least squares solution in Equation (5).

#### 2.2.2 Effectiveness of standard aggregation

To understand what is lost when aggregating a model, it is helpful to quantify the error introduced by representing the larger model with a reduced one. This aggregation error is defined by 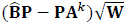, which is an *m* × *n* matrix of residuals from the weighted least squares fit (predicted – actual). To gauge the fit of the aggregated model we use the effectiveness of standard aggregation (after Iwasa et al., 1989),

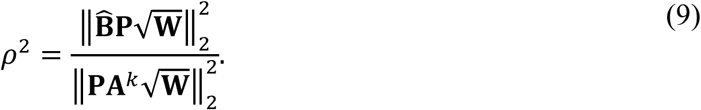

The effectiveness of standard aggregation is analogous to the coefficient of determination in linear regression. When there is perfect aggregation, **B̂P** = **PA**^*k*^, and so *ρ*^2^ = 1; otherwise, *ρ*^2^ < 1 (Hinrichsen, 2023).

#### 2.2.3 Properties of the standard aggregated matrix

The standard aggregated matrix **B̂** inherits the following properties from the original matrix: it is a Leslie matrix, it is irreducible whenever **A** is irreducible, and it is primitive (irreducible with a strictly dominant Perron eigenvalue) whenever **A** is primitive, its population growth rate is *λ*^*k*^, and its stable age distribution is **Pw** (Hinrichsen, 2023). These last two properties demonstrate that the population growth rate and stable age distribution of **B̂** are both consistent with the original model.

The standard aggregated matrix has reproductive values that are inconsistent with those of **A**, which, in turn, leads to inconsistent elasticities—a key limitation to comparative demography (Salguero-Gómez & Plotkin, 2010). Reproductive values of the aggregated matrix **B** are consistent when they are **v**_**B**_ = (**PWP**^⊤^)^−1^**PWv**, yielding weighted averages of the original reproductive values based on the stable age distribution (e.g. for *k* = 2, the reproductive value for the first aggregated age class is (*w*_1_*v*_1_ + *w*_2_*v*_2_)/(*w*_1_ + *w*_2_)). Elasticities are consistent when E(**B**) = **P**E(**A**^*k*^)**P**^⊤^, where E(·) returns the elasticity matrix of *λ*^*k*^ with respect to the entries of the indicated matrix. Elasticities are taken with respect to the entries of **A**^*k*^, since aggregation is applied to **A**^*k*^ rather than **A**.

We illustrate the inconsistency of the standard aggregator using a Leslie matrix from the COMADRE Animal Matrix Database (ID 248204; accessed 12 January 2026, version 4.23.3.1), corresponding to a population of common roach (*Rutilus rutilus*) (Otjacques et al., 2016; Salguero-Gómez et al., 2016a). Figure 2 shows that using the standard aggregator to reduce the 10 × 10 matrix **A**^2^ to a 5 × 5 matrix **B̂** yields inconsistent reproductive values and elasticities.

**FIGURE 2.**
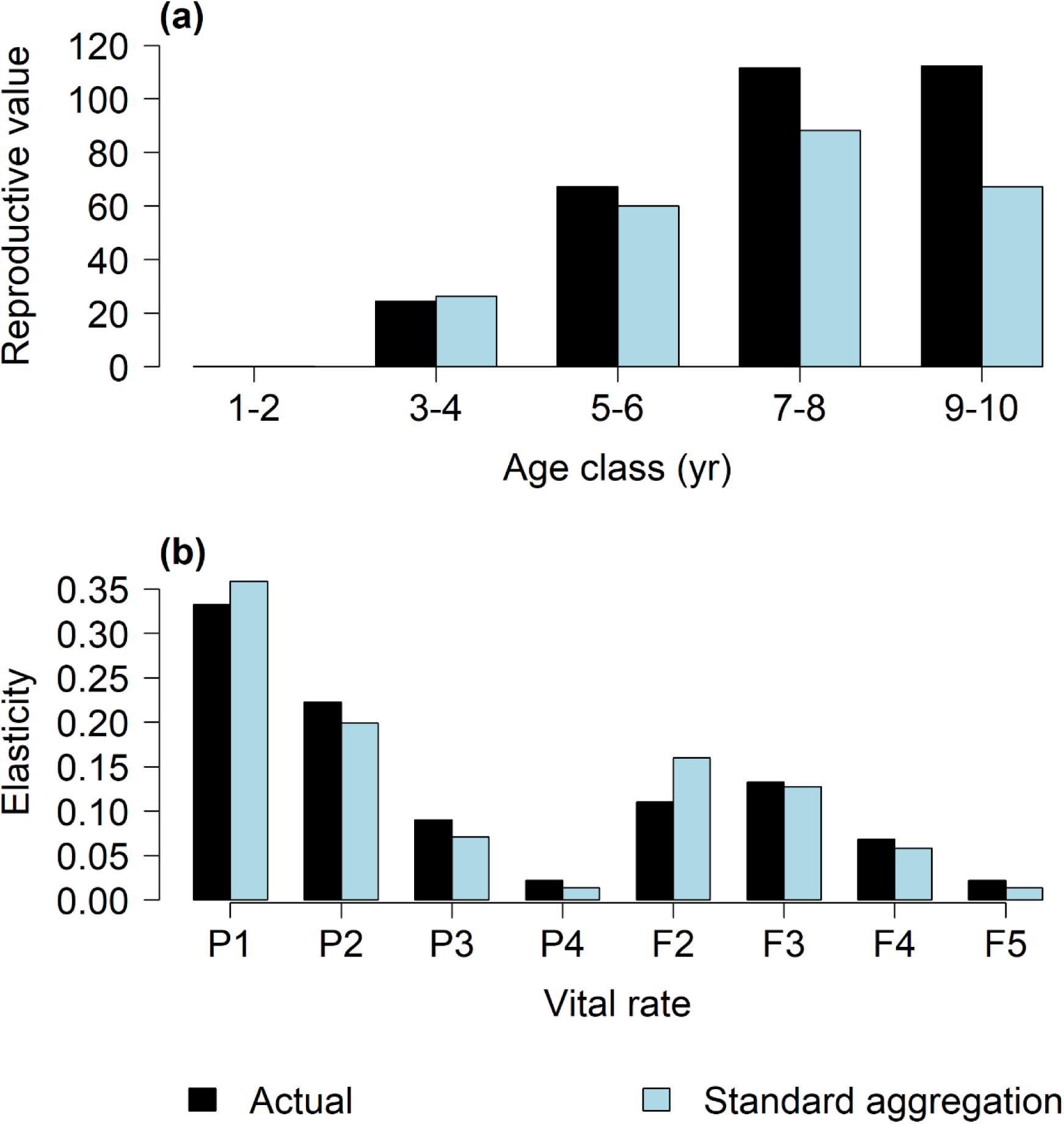
The standard aggregator yields inconsistent reproductive values and elasticities. Illustration showing the inconsistency of (a) reproductive values and (b) elasticities of population growth rate (*λ*) to vital rates, resulting from standard aggregation of a matrix population model (MPM) for common roach (*Rutilus rutilus*). The original MPM has dimensionality *n* = 10, and the aggregated MPM has dimensionality *m* = 5. ‘Actual’ values are those expected if aggregation preserved consistency with the original MPM. Vital rates of the aggregated Leslie MPM are: *P*_*i*_ (survival probability) and *F*_*i*_ (fertility rate) for aggregated age class *i*.

### 2.3 An elasticity-consistent aggregator

To derive the elasticity-consistent aggregator, we apply the method of interstage flows to the balanced matrix **Ã** = **VAV**^−1^, where **V** = diag(**v**) (Appendix A); this transformation yields scale-invariant interstage flows (Hinrichsen, 2025; Yokomizo et al., 2025). The resulting elasticity-consistent aggregated matrix is

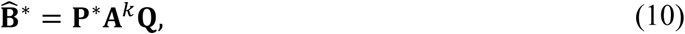

(where **P**^∗^ = **PWP**^⊤^(**PVWP**^⊤^)^−1^**PV** (an *m* × *n* matrix) and **Q** is given by Equation (6). The matrix **B̂**^∗^ has stable population growth rate, stable age distribution, reproductive values, and elasticities that are all consistent with the original model (Appendix A). This method agrees with Bienvenu et al. (2017, Eq. 19) who aggregated a general MPM, and used a more general partitioning matrix, which can lead to survival probabilities > 1. We will show that our choice to apply the method to Leslie MPMs and to aggregate **A**^*k*^ with partitioning matrix **P**, guarantees that **B̂**^∗^ is a Leslie matrix with survival probabilities ≤ 1.

#### 2.3.1 Effectiveness of elasticity-consistent aggregation

The effectiveness of elasticity-consistent aggregation is given by Equation (9) applied to balanced matrices:

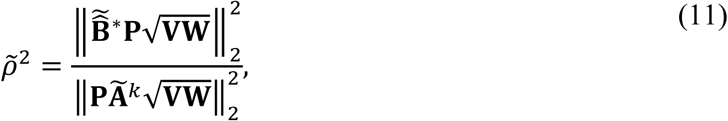

where **B̂̃**^∗^ = **PÃ**^*k*^**VWP**^⊤^(**PVWP**^⊤^)^−1^ is the balanced form of **B̂**^∗^ and **Ã**^*k*^ = **VA**^*k*^**V**^−1^ is the balanced form of **A**^*k*^, and the aggregation error is the *m* × *n* matrix (**B̂̃**^∗^**P** − **PÃ**^*k*^). The metric *ρ̃*^2^ never exceeds 1 and perfect aggregation occurs precisely when *ρ̃*^2^ = 1. There is perfect aggregation (i.e. *ρ̃*^2^ = 1) whenever *m* = 1 (Appendix B).

#### 2.3.2 Properties of the elasticity-consistent aggregated matrix

The elasticity-consistent aggregated matrix preserves stable growth rate, stable age distribution, and reproductive values (Appendix A). In addition, it is a Leslie matrix and is irreducible or primitive whenever **A** is irreducible or primitive (Appendix C). Because **B̂**^∗^and **B̂** share identical survival probabilities, they can differ only in their fertility rates.

#### 2.3.3 When *n* is not a multiple of *m*

When the dimensionality of the original matrix is not an integer multiple of that of the aggregated matrix (i.e. *n* = *km*, where *k* is a non-integer), we follow Hinrichsen (2023) by first disaggregating the original MPM into a larger MPM of dimensionality *nm* that preserves the original dynamics, and then aggregating this disaggregated model (Appendix D). The resulting disaggregated matrix is imprimitive, that is, irreducible with multiple eigenvalues of maximum absolute value.

Applying the standard aggregator to the disaggregated matrix preserves the following characteristics of the original MPM: Leslie form, irreducibility, primitivity, and consistency of stable growth rate and stable age distribution (Hinrichsen, 2023). The elasticity-consistent aggregator preserves these properties and additionally yields reproductive values consistent with the original MPM (Appendix D).

## 3. APPLICATION TO TWELVE ANIMAL POPULATIONS

### 3.1 Demographic data

To compare the accuracies of the standard and elasticity-consistent aggregators, we applied both to MPMs selected from the COMADRE Animal Matrix Database (accessed 12 January 2026, version 4.23.3.1). We accessed COMADRE through the R package ‘Rcompadre’ (version 1.4.0, Jones et al., 2022), which uses the R statistical language and environment (version 4.5.2, R Core Team, 2025). To ensure broad phylogenetic and life-history coverage, we selected 12 age-structured MPMs, each from a different taxonomic class. We chose MPMs whose growth rates, dimensionalities, and projection intervals spanned the interquartile ranges of these traits across all COMADRE Leslie matrices and included both primitive and imprimitive forms. As an exception, all selected MPMs had dimensionality >3 to allow at least three aggregated MPMs.

Details of the selected MPMs and associated species are presented in Table S1. Growth rates ranged from 0.47 (earthworm) to 2.39 (soft-shell clam) and dimensionalities from 4 (axolotl) to 50 (freshwater crocodile) across the selected MPMs. Projection intervals ranged from 0.01 year (water flea) to 1 year (e.g. freshwater crocodile). One of the MPMs had an imprimitive population projection matrix (bog fritillary), giving rise to persistent oscillations with a period of five years. To ensure that reproductive values were positive, we dropped the postreproductive age classes of the Rocky Mountain elk and amphipod populations, making their associated matrices irreducible (Leslie, 1945). We transformed the original Lefkovitch matrices for soft-shell clam and earthworm into Leslie matrices of dimensionality 11 and 7, respectively (Appendix E). The rest of the MPMs were of Leslie form.

### 3.2 Aggregation

We aggregated the Leslie MPMs using both standard and elasticity-consistent methods. If *m* did not divide *n* evenly, we used a disaggregated MPM in place of the original MPM to construct the aggregated MPM of dimensionality *m* (see Section 2.3.3). When *m* = *n*, the ‘aggregated’ MPM was the original MPM.

### 3.3 Comparing accuracies of aggregators using demographic parameters

We compared standard and elasticity-consistent aggregation using fertility-dependent demographic parameters, generation time, Demetrius’ entropy, and net reproductive rate, because both methods yield identical survival probabilities. Accuracy was evaluated across aggregation levels by comparing estimates from aggregated matrices to the corresponding values from the original Leslie MPMs. Performance was assessed using three metrics: a win rate, scored as 1, ½, or 0 when the elasticity-consistent estimate was closer to, equally close to, or further from the original value than the standard estimate; median absolute error (MAE) for each method; and directional error, expressed as the proportions of estimates that were above (too high), below (too low), or equal to (exact) the original value. Total win rates and directional errors were computed by pooling comparisons across all aggregation levels and MPMs.

#### 3.3.1 Generation time

First, we compared estimates of generation time, a strong predictor of the location of a species’ population along the slow-fast continuum (Gaillard et al., 2005). This metric is also an excellent proxy to examine the temporal scales at which populations respond to perturbations and natural selection (Brown et al., 2022; Ellner, 2018), and when available is included in the IUCN Red List status criteria (Cooke et al., 2018). We used the generation time of Bienvenu and Legendre (2015),

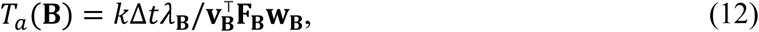

where *k*Δ*t* is the projection interval of the aggregated model (in years), *λ*_**B**_ is the stable growth rate of the aggregated matrix **B**, **w**_**B**_is the stable age distribution of the aggregated matrix **B**, **F**_**B**_is the fertility matrix of the aggregated matrix **B** with fertility rates in the first row and 0s elsewhere, and **v**_**B**_is the vector of reproductive values of **B** such that **v**^⊤^**w**_**B**_ = 1. When **B** = **A**, the generation time is *T*_*a*_(**A**) = Δ*t*λ/**v**^⊤^**Fw**, where **F** is the fertility matrix of **A**.

#### 3.3.2 Demetrius’ entropy

Second, we compared estimates of Demetrius’ entropy,

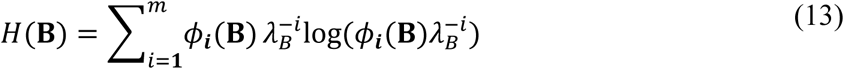

where *ɸ*_***i***_(**B**) are derived from the characteristic equation for 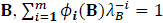 (Caswell, 2001). Demetrius’ entropy measures the spread of reproduction through the life cycle. It equals zero precisely when reproduction occurs in a single age class and is maximal precisely when reproduction is evenly distributed across ages (Caswell, 2001; Salguero-Gómez et al., 2016b).

#### 3.3.3 Net reproductive rate

Finally, we compared estimates of net reproductive rate,

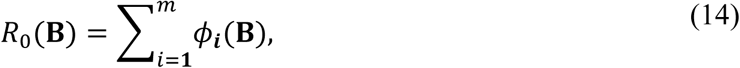

which represents the expected number of offspring produced by an individual over its lifetime (Caswell, 2001).

### 3.4 Effectiveness of aggregation

As an additional test of accuracy, we evaluated the effectiveness of aggregation measures *ρ*^2^ and *ρ̃*^2^ against the dimensionality of the aggregated MPMs. The metric *ρ̃*^2^ applies to balanced matrices, where age classes are weighted by reproductive values. Comparing *ρ*^2^and *ρ̃*^2^ reveals how balancing affects aggregation error.

## 4. RESULTS

### 4.1 Demographic parameter estimates

#### 4.1.1 Generation time

The generation time estimates are given in Figure 3 and the win rate, MAE, and directional errors are tabulated in Table S2. Overall, elasticity-consistent estimates were more accurate than the standard estimates, with a win rate of 86%. Directional error showed that 24% of the elasticity-consistent estimates were too high, 0% were too low, and 76% were exact. For the standard estimates, 20% were too high, 77% were too low, and 3% were exact. Regardless of the aggregator, there was no error in the estimates for bog fritillary (imprimitive) and little error for the amphipod matrix (nearly imprimitive).

**FIGURE 3.**
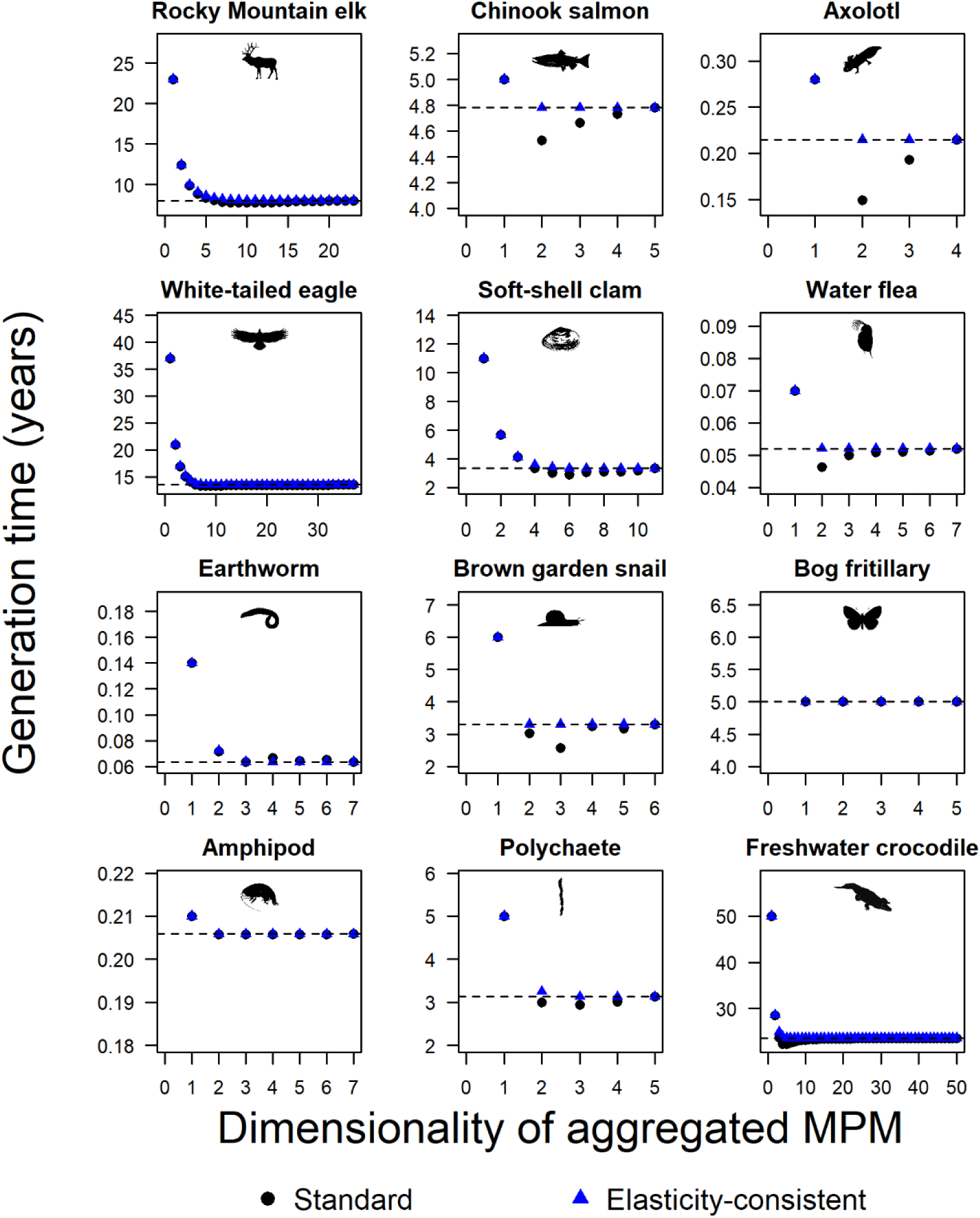
Generation time is often more accurately estimated by the elasticity-consistent aggregator across 12 animal species. Depicted is generation time estimated from standard aggregator and elasticity-consistent aggregator, both evaluated against the dimensionality of the aggregated matrix population model (MPM), *m*. The dashed horizontal line represents the actual generation time, derived from the original (unaggregated) MPM. The silhouettes are from PhyloPic (https://www.phylopic.org; Keesey, 2023) and were added using the rphylopic R package (Gearty & Jones, 2023). Silhouettes were contributed as follows: *Cervus canadensis* by Richard Rich (2025; CC0 1.0); *Oncorhynchus tshawytscha* by Xavier Giroux-Bougard (2018; PDM 1.0); *Ambystoma mexicanum* by Chuanixn Yu (2021; CC0 1.0); *Haliaeetus albicilla* by Andy Wilson (2023; CC0 1.0); *Mya arenaria* by Marina Vingiani (2025; CC BY 4.0); *Daphnia* by T. Michael Keesey (2013; PDM 1.0); *Eiseniella tetraedra* by Nathan Jay Baker (2024; CC0 1.0); *Helix aspersa* by Yan Wong from photo by Denes Emoke (2016; CC0 1.0); *Boloria eunomia* by Didier Descouens (2025; CC BY 4.0); *Ampelisca agassizi* by Collin Gross (2021; CC BY 3.0); *Nephtys* by Guillaume Dera (2023; CC0 1.0); *Crocodylus* by Becky Barnes (2021; CC0 1.0).

#### 4.1.2 Demetrius’ entropy

The Demetrius’ entropy estimates are given in Figure S1 and win rate, MAE, and directional errors are tabulated in Table S3. Overall, the elasticity-consistent estimates of Demetrius’ entropy were slightly more accurate than the standard estimates, with a win rate of 60%. However, the win rates and MAEs indicated that the standard estimates were more accurate for the Chinook salmon, axolotl, earthworm, and amphipod MPMs (Table S3). Directional error showed that 7% of the elasticity-consistent estimates were too high, 90% were too low, and 3% were exact. For the standard estimates, 6% were too high, 91% were too low, and 3% were exact.

#### 4.1.3 Net reproductive rate

The net reproductive rate estimates are given in Figure S2 and win rate, MAE, and directional errors are tabulated in Table S4. Although the stable growth rates are guaranteed to be exactly consistent with both methods of aggregation, its lifetime reproductive counterpart, net reproductive rate, is not (Figure S2). Overall, the elasticity-consistent estimates of net reproductive rate were more accurate than the standard estimates, with a win rate of 76%. However, win rates and MAEs indicate that the standard estimates were more accurate for the softshell clam, water flea, and earthworm MPMs (Table S4). Directional error showed that 90% of the elasticity-consistent estimates were too high, 7% were too low, and 3% were exact. For the standard estimates, 39% were too high, 58% were too low, and 3% were exact.

### 4.2 Effectiveness of aggregation

Effectiveness of elasticity-consistent aggregation was higher than effectiveness of standard aggregation (Figure 4). Thus, weighting age classes by their reproductive values reduces aggregation error. The plots confirmed that *ρ̃*^2^ = 1 (i.e. perfect aggregation with balancing) whenever *m* = 1 (Appendix B).

**FIGURE 4.**
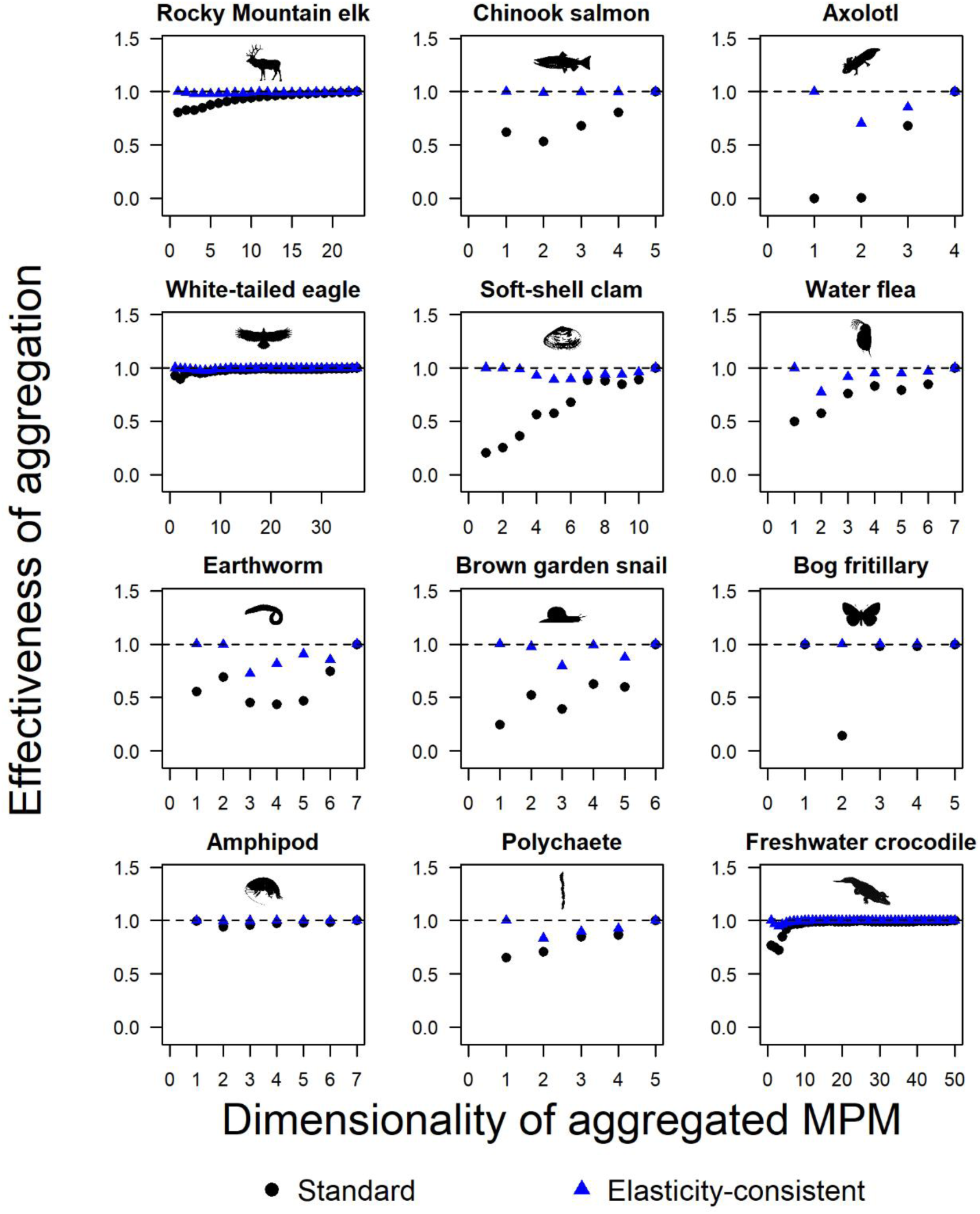
Effectiveness of aggregation is higher using elasticity-consistent aggregation across 12 animal species. Depicted is the effectiveness of aggregation for the standard aggregator and elasticity-consistent aggregator, evaluated against the dimensionality of the aggregated matrix population model (MPM), *m*. The dashed horizontal line represents an effectiveness of one, corresponding to perfect aggregation (no aggregation error). Silhouettes are identical to those in Figure 3; credits are given in the Figure 3 caption.

Balancing by reproductive values increases aggregation effectiveness because balanced matrix entries are less variable and therefore easier to fit by aggregation. Figure S3a compares predicted values 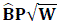 with actual values 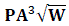 for the unbalanced brown garden snail matrix (*n* = 6, *m* = 3). Figure S3b shows the corresponding comparison for balanced matrices, with predicted values 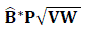 and actual values 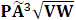. Balancing lowers transitions into the youngest age class and increases transitions into older classes according to their reproductive values.

Imprimitive matrices show the lowest aggregation error. When reproduction is confined to the oldest age class, as in the bog fritillary, reduced models capture the dynamics exactly (Figure 4), a pattern also seen in the nearly imprimitive amphipod matrix. However, aggregation error depends on the method: under the standard aggregator, the bog fritillary exhibits high error (*ρ*^2^ = 0.14) at *m* = 2, whereas the elasticity-consistent aggregator showed no aggregation error (*ρ̃*^2^ = 1) across all *m*.

## DISCUSSION

As comparative demography increasingly draws on large, heterogeneous demographic databases, the challenge towards identifying generality in ecology lies in making biologically meaningful inferences from matrix population models (MPMs) that differ widely in complexity (Che-Castaldo et al., 2020; Gascoigne et al., 2023; Kendall, 2015; Römer et al., 2024; Salguero-Gómez et al., 2021). The growth of large, open-access datasets, which enables the discovery of broad life-history patterns across taxa, also magnifies the challenges of author-driven choices in model formulation. The ability of comparative demography to discover generalities depends on whether observed patterns faithfully reflect underlying biological processes rather than artifacts of model formulation.

To compare two MPMs of different complexity, one can aggregate the more complex MPM so that its classes coincide with those of the smaller MPM (Enright et al., 1995; Salguero-Gómez & Plotkin, 2010). Here we evaluate the accuracy of two such aggregators, the standard and elasticity consistent aggregators, and show that they differ fundamentally in their ability to preserve key demographic properties of Leslie MPMs during dimensionality reduction. In our analysis across 12 diverse animal species, generation time, Demetrius’ entropy, and net reproductive rate were more accurately estimated by the elasticity-consistent approach in 86%, 60%, and 76% of cases, respectively.

There are clear advantages of elasticity-consistent aggregation methods for comparative demography. Variation in model complexity has hindered comparisons in databases COMADRE and COMPADRE (Salguero-Gómez et al., 2015, 2016a), because we lack the necessary tools to make unbiased comparisons between MPMs that differ in construction (Kendall et al., 2015; Salguero-Gómez & Plotkin, 2010). By providing a rigorous way to reduce complex models while preserving key demographic measures, the elasticity-consistent aggregator offers a unifying framework for life-history synthesis that enables researchers to make fairer comparisons across diverse models in analyses of elasticity, life-history trade-offs, and the slow–fast continuum (Gaillard et al., 1989; Stott et al., 2024).

Although Bienvenu et al. (2017) independently developed an elasticity-preserving aggregator, our approach differs in derivation, implementation, and scope. First, we use balancing and the method of interstage flows as a shortcut to weighted least squares to derive the elasticity-consistent aggregator while Bienvenu et al. (2017) use a Markov chain representation of the MPM. Second, we aggregate Leslie MPM → Leslie MPM while Bienvenu et al. (2017) aggregate general MPM → general MPM, which can produce survival probabilities > 1. Third, in contrast to Bienvenu et al. (2017), our derivation does not require primitivity of the original population projection matrix. In fact, we found that imprimitive population projection matrices harbour the greatest potential for accurate aggregation (e.g. bog fritillary MPM) and are used in the disaggregation step that allows aggregation of a Leslie MPM → Leslie MPM when *m* does not divide *n*. Fourth, we incorporate explicit aggregation-effectiveness metrics (*ρ*^2^ and *ρ̃*^2^), providing quantitative diagnostics of aggregation error absent in Bienvenu et al. (2017).

Our approach highlights the central role of interstage flows (Kawano et al., 1987; Yokomizo et al., 2024) in aggregation. By weighting by stable age distribution, interstage flow is the currency that allows vital rates to be summed across age classes. Without such weighting, vital rates cannot be safely summed because their associated age classes differ in natural abundance. Further balancing by reproductive values makes these interstage flows scale invariant (Hinrichsen, 2025), which leads to an aggregator that not only preserves reproductive values and elasticities, but commutes with change of units (that is, changing units, aggregating, and changing back to the original units gives the same MPM as aggregating alone) (Bienvenu et al., 2017).

A key challenge of aggregation is that it can obscure demographic and dynamic details by ignoring individual heterogeneity within groups. This limitation motivates the use of a disaggregator, which preserves demographic detail by expanding smaller MPMs to match the structure of larger MPMs. Because disaggregation is a one-to-many operation, yielding infinitely many disaggregated MPMs consistent with a MPM, resolving the ambiguity of this inverse problem may require a Bayesian approach that incorporates prior demographic knowledge (Clark, 2005; Tarantola, 2005). Integral projection models carry disaggregation to the extreme, making discrete age classes continuous (Easterling et al., 2000). In our framework, disaggregation is not merely a theoretical inverse of aggregation; it allows aggregation when the dimensionalities of the original and aggregated Leslie MPMs are incompatible (see Section 2.3.3).

Although we have focused on aggregation of Leslie MPMs, standard and elasticity-consistent aggregators may be extended to size- or stage-structured MPMs (Bienvenu et al., 2017; Salguero-Gómez & Plotkin, 2010), and multiregional MPMs (Auger et al., 2008). In the general case, there is freedom to choose what stages (with age as a special case) to combine by applying an arbitrary partitioning matrix: one is not restricted to combining adjacent stages and some stages may not be combined at all (Bienvenu et al., 2017; Hooley, 2000; Salguero-Gómez & Plotkin, 2010). If we relax the Leslie MPM → Leslie MPM requirement and apply aggregators directly to a general MPM without changing the projection interval, the elasticity-consistent aggregator preserves mean generation time for any partition (Bienvenu et al., 2017). Yearsley & Fletcher (2002) also provide an aggregator that preserves generation time, though not elasticities. When comparing general MPMs, it is a challenge to align stages so that they are comparable (Salguero-Gómez & Plotkin, 2010). In contrast, aligning age classes of Leslie MPMs is relatively straightforward (Hinrichsen, 2023), although one must consider census type (Caswell, 2001). Furthermore, it is an unsolved problem aggregating general MPM → general MPM such that elasticities are preserved and aggregated survival probabilities ≤ 1 (Bienvenu et al., 2017). Applying aggregation to Leslie MPMs is less restrictive than it may first appear because stage-classified MPMs can be cast as Leslie MPMs using age-from-stage methods (Caswell, 2001; Cochran & Ellner, 1992).

We conclude that the elasticity-consistent aggregator, by preserving key demographic properties of the original MPM, provides a robust framework to compare MPMs of varying complexity drawn from big demographic data sets. This approach advances comparative demography, and warrants further exploration across a wider spectrum of MPMs to help seize the promise of big data in ecology and discover principles across the tree of life, from microbes to fungi, plants, and animals.

## Acknowledgements

R.A.H. is grateful to Charlie Johnson for suggesting Leslie model aggregation as a line of research while he was a graduate student at Clemson University and to Eric Howe, Yoh Iwasa, and Simon Levin for discussions. R.S.G. was supported by a NERC Pushing the Frontiers grant (NE/X013766/1). H.Y. was supported by JSPS KAKENHI Grant Number 25K09778.

## Data availability

Data available via https://compadre-db.org/Data/Compadre (Salguero-Gómez et al., 2015).

## Author contributions

R.A.H. developed the aggregation methods, linking them to H.Y.’s interstage flow matrix and its scale-invariant extension. R.S.-G. conceptualized the application of aggregation theory to comparative demography and revealed the limitations of conventional aggregators. R.A.H., with assistance from H.Y., implemented the software and wrote the R code for analyses and figure generation. R.A.H. led the writing, and all authors contributed to the manuscript drafts and gave final approval for publication.

