## Appendices for "From theory to application: Elasticity-consistent aggregation of Leslie matrix population models for comparative demography"

### APPENDIX A Elasticity-consistent aggregator

#### A.1 Derivation

We now derive an elasticity-consistent aggregator by insisting that both the stable age distribution and the reproductive values are consistent with those of $\mathbf{A}$. To do this, we apply the method of interstage flows to balanced forms of the matrices. To balance $\mathbf{A}$, we rescale the matrix population model by reproductive values, which yields the balanced matrix

$$\widetilde{\mathbf{A}} = \mathbf{V}\mathbf{A}\mathbf{V}^{-1}, \tag{A.1}$$

where $\mathbf{V} = \mathrm{diag}(\mathbf{v})$. This balanced matrix governs the dynamics of the age-specific total reproductive values, $\mathbf{v} \circ \mathbf{x}(t)$ and it has stable age distribution $\widetilde{\mathbf{w}} = \mathbf{v} \circ \mathbf{w}$ and reproductive values of $\tilde{\mathbf{v}} = \mathbf{e}$ where $\mathbf{e}$ is a vector of 1s (Hinrichsen, 2025). We call $\widetilde{\mathbf{A}}$ a 'balanced' matrix, which denotes a matrix derived via a diagonal similarity transformation (Golub & Van Loan, 2013). Balancing prevents undue influence of immature classes and makes the interstage flows scale invariant (Hinrichsen, 2025; Yokomizo et al., 2025). Most importantly, for the sake of aggregation, balancing expresses age classes in a common currency that allows them to be added together with strict regard to their reproductive values.

The balanced form of $\mathbf{A}^k$ is $\widetilde{\mathbf{A}}^k = \mathbf{V}\mathbf{A}^k\mathbf{V}^{-1}$, which, like $\widetilde{\mathbf{A}}$, has stable age distribution $\widetilde{\mathbf{w}} = \mathbf{v} \circ \mathbf{w}$ and reproductive values of $\tilde{\mathbf{v}} = \mathbf{e}$, where $\mathbf{e}$ is a vector of 1s. Let $\widehat{\mathbf{B}}^*$ denote the new elasticity-consistent aggregated matrix. For consistency, the stable age distribution of $\widehat{\mathbf{B}}^*$ must be $\mathbf{w}_{\widehat{\mathbf{B}}^*} = \mathbf{P}\mathbf{w}$, its reproductive values must be $\mathbf{v}_{\widehat{\mathbf{B}}^*} = (\mathbf{P}\mathbf{W}\mathbf{P}^\top)^{-1}\mathbf{P}\mathbf{W}\mathbf{v}$, and $\lambda\big(\widehat{\mathbf{B}}^*\big) = \lambda(\mathbf{A}^k) = \lambda^k$.

We derive $\widehat{\mathbf{B}}^*$by the method of interstage flows, which equates the interstage flows of the balanced form of $\widehat{\mathbf{B}}^*$ with the corresponding interstage flows of the balanced form of $\mathbf{A}^k$ as follows:

$$\mathcal{F}\left(\widetilde{\widehat{\mathbf{B}}}{}^*\right) = \mathbf{P}\mathcal{F}\big(\widetilde{\mathbf{A}}^k\big)\mathbf{P}^\top. \qquad \text{(A.2)}$$

The interstage flow matrix of $\widetilde{\mathbf{A}}^k$ is

$$\begin{aligned}\mathcal{F}\big(\widetilde{\mathbf{A}}^k\big) &= (\mathbf{V}\mathbf{A}^k\mathbf{V}^{-1})\mathbf{V}\mathbf{W} \\ &= \mathbf{V}\mathbf{A}^k\mathbf{W}\end{aligned} \qquad \text{(A.3)}$$

Let $\mathbf{W}_{\widehat{\mathbf{B}}^*} = \mathrm{diag}(\mathbf{w}_{\widehat{\mathbf{B}}^*}) = \mathbf{PWP}^\top$, and let $\mathbf{V}_{\widehat{\mathbf{B}}^*} = \mathrm{diag}(\mathbf{v}_{\widehat{\mathbf{B}}^*}) = \mathbf{PVWP}^\top(\mathbf{PWP}^\top)^{-1}$ . Then, the interstage flow of $\widetilde{\widehat{\mathbf{B}}}^*$ **is**

$$\begin{aligned}\mathcal{F}\left(\widetilde{\widehat{\mathbf{B}}}^*\right) &= (\mathbf{V}_{\widehat{\mathbf{B}}^*}\widehat{\mathbf{B}}^*\mathbf{V}_{\widehat{\mathbf{B}}^*}{}^{-1})\mathbf{V}_{\widehat{\mathbf{B}}^*}\mathbf{W}_{\widehat{\mathbf{B}}^*} \\ &= \mathbf{V}_{\widehat{\mathbf{B}}^*}\widehat{\mathbf{B}}^*\mathbf{W}_{\widehat{\mathbf{B}}^*} \\ &= \mathbf{PVWP}^\top(\mathbf{PWP}^\top)^{-1}\widehat{\mathbf{B}}^*\mathbf{PWP}^\top.\end{aligned} \tag{A.4}$$

Substituting these expressions for the interstage flow matrices in Equation (A.2) produces

$$(\mathbf{PVWP}^\top)(\mathbf{PWP}^\top)^{-1}\widehat{\mathbf{B}}^*(\mathbf{PWP}^\top) = \mathbf{PVA}^k\mathbf{WP}^\top. \tag{A.5}$$

Solving for $\widehat{\mathbf{B}}^*$ in Equation (A.5) yields

$$\begin{aligned}\widehat{\mathbf{B}}^* &= \mathbf{PWP}^\top(\mathbf{PVWP}^\top)^{-1}\mathbf{PVA}^k\mathbf{WP}^\top(\mathbf{PWP}^\top)^{-1} \\ &= \mathbf{P}^*\mathbf{A}^k\mathbf{Q},\end{aligned} \tag{A.6}$$

where $\mathbf{P}^* = \mathbf{PWP}^\top(\mathbf{PVWP}^\top)^{-1}\mathbf{PV}$ and $\mathbf{Q} = \mathbf{WP}^\top(\mathbf{PWP}^\top)^{-1}$ .

#### A.2 Consistency of asymptotic dynamics

Here we show that $\widehat{\mathbf{B}}^*$ has stable growth rate, stable age distribution, and reproductive values that are all consistent with the original matrix $\mathbf{A}$, that is: $\lambda(\widehat{\mathbf{B}}^*) = \lambda(\mathbf{A}^k)$, $\mathbf{w}_{\widehat{\mathbf{B}}^*} = \mathbf{Pw}$, and $\mathbf{v}_{\widehat{\mathbf{B}}^*} = (\mathbf{PWP}^\top)^{-1}\mathbf{PWv}$. We first show that $\mathbf{w}_{\widehat{\mathbf{B}}^*} = \mathbf{Pw}$ is the stable age distribution because it is a right eigenvector corresponding to dominant eigenvalue $\lambda(\widehat{\mathbf{B}}^*) = \lambda(\mathbf{A}^k)$.

$$\begin{aligned}
\widehat{\mathbf{B}}^*\mathbf{Pw} &= \mathbf{PWP}^\top(\mathbf{PVWP}^\top)^{-1}\mathbf{PVA}^k\mathbf{WP}^\top(\mathbf{PWP}^\top)^{-1}\mathbf{Pw} \qquad \text{(A.7)}\\
&= \mathrm{diag}(\mathbf{Pw})\left\{\mathrm{diag}\big(\mathbf{P}(\mathbf{v}\circ\mathbf{w})\big)\right\}^{-1}\mathbf{P}\mathrm{diag}(\mathbf{v})\mathbf{A}^k\mathrm{diag}(\mathbf{w})\mathbf{P}^\top\{\mathrm{diag}(\mathbf{Pw})\}^{-1}\mathbf{Pw}\\
&= \mathrm{diag}(\mathbf{Pw})\left\{\mathrm{diag}\big(\mathbf{P}(\mathbf{v}\circ\mathbf{w})\big)\right\}^{-1}\mathbf{P}\mathrm{diag}(\mathbf{v})\mathbf{A}^k\mathrm{diag}(\mathbf{w})\mathbf{P}^\top\mathbf{e}\\
&= \mathrm{diag}(\mathbf{Pw})\left\{\mathrm{diag}\big(\mathbf{P}(\mathbf{v}\circ\mathbf{w})\big)\right\}^{-1}\mathbf{P}\mathrm{diag}(\mathbf{v})\mathbf{A}^k\mathbf{w}\\
&= \mathrm{diag}(\mathbf{Pw})\left\{\mathrm{diag}\big(\mathbf{P}(\mathbf{v}\circ\mathbf{w})\big)\right\}^{-1}\mathbf{P}(\mathbf{v}\circ\mathbf{w})\lambda(\mathbf{A}^k)\\
&= \mathrm{diag}(\mathbf{Pw})\mathbf{e}\lambda(\mathbf{A}^k)\\
&= \lambda(\mathbf{A}^k)\mathbf{Pw},
\end{aligned}$$

where $\mathbf{e}$ is an $m$-vector of 1s. Equation (A.7) shows that, indeed, $\mathbf{Pw}$ is the stable age distribution of $\widehat{\mathbf{B}}^*$ corresponding to dominant eigenvalue $\lambda\left(\widehat{\mathbf{B}}^*\right) = \lambda(\mathbf{A}^k)$.

Next, we show consistency of reproductive values, that is $\mathbf{v}_{\widehat{\mathbf{B}}^*} = (\mathbf{PWP}^\top)^{-1}\mathbf{PWv}$ is a left eigenvector of $\widehat{\boldsymbol{B}}^*$ associated with dominant eigenvalue $\lambda\left(\widehat{\mathbf{B}}^*\right) = \lambda(\mathbf{A}^k)$.

$$
\begin{aligned}
\mathbf{v}^\top\mathbf{WP}^\top(\mathbf{PWP}^\top)^{-1}\widehat{\boldsymbol{B}}^* &= \mathbf{v}^\top\mathbf{WP}^\top(\mathbf{PWP}^\top)^{-1}\mathbf{PWP}^\top(\mathbf{PVWP}^\top)^{-1}\mathbf{PVA}^k\mathbf{WP}^\top(\mathbf{PWP}^\top)^{-1} \\
&= \mathbf{v}^\top\mathbf{WP}^\top(\mathbf{PVWP}^\top)^{-1}\mathbf{PVA}^k\mathbf{WP}^\top(\mathbf{PWP}^\top)^{-1} \\
&= (\mathbf{v}\circ\mathbf{w})^\top\,\mathbf{P}^\top\left\{\mathrm{diag}\big(\mathbf{P}(\mathbf{v}\circ\mathbf{w})\big)\right\}^{-1}\mathbf{P}\mathrm{diag}(\mathbf{v})\mathbf{A}^k\mathbf{WP}^\top(\mathbf{PWP}^\top)^{-1} \\
&= \mathbf{e}^\top\mathbf{P}\mathrm{diag}(\mathbf{v})\mathbf{A}^k\mathbf{WP}^\top(\mathbf{PWP}^\top)^{-1} \\
&= \mathbf{v}^\top\mathbf{A}^k\mathbf{WP}^\top(\mathbf{PWP}^\top)^{-1} \\
&= \lambda(\mathbf{A}^k)\mathbf{v}^\top\mathbf{WP}^\top(\mathbf{PWP}^\top)^{-1}.
\end{aligned}
\tag{A.8}
$$

Equation (A.8) shows that, indeed, $\mathbf{v}_{\widehat{\mathbf{B}}^*} = (\mathbf{PWP}^\top)^{-1}\mathbf{PWv}$ is the dominant left eigenvector of $\widehat{\boldsymbol{B}}^*$ corresponding to dominant eigenvalue $\lambda\left(\widehat{\mathbf{B}}^*\right) = \lambda(\mathbf{A}^k)$.

**A.3 Consistency of elasticities**

Consistency of elasticities follows because the elasticity matrix is the normalized interstage flow of a population projection matrix balanced by its reproductive values (Hinrichsen, 2025). Therefore,

$$
\begin{aligned}
\mathrm{E}\left(\widehat{\mathbf{B}}^{*}\right) &= \mathcal{F}\left(\widetilde{\widehat{\mathbf{B}}}^{*}\right)/\lambda^{k} \qquad \text{(A.9)} \\
&= \mathbf{P}\mathcal{F}(\widetilde{\mathbf{A}}^{k})\,\mathbf{P}^{\top}/\lambda^{k} \\
&= \mathbf{P}\mathcal{F}(\mathbf{A}^{k})\mathbf{P}^{\top}.
\end{aligned}
$$

The second line of Equation (A.9) follows from Equation (A.2).

**APPENDIX B Effectiveness of elasticity-consistent aggregation is 1 when the dimensionality of the aggregated matrix is 1**

In this appendix, we show that the effectiveness of elasticity-consistent aggregation, $\tilde{\rho}^2$, is 1 (perfect aggregation) when $m = 1$. Recall that $\mathbf{V} = \mathrm{diag}(\mathbf{v})$, $\mathbf{W} = \mathrm{diag}(\mathbf{w})$, where $\mathbf{v}$ is the vector of reproductive values of $\mathbf{A}$ and $\mathbf{w}$ is the stable age distribution of $\mathbf{A}$, and the dominant eigenvalue of $\mathbf{A}$ is $\lambda$. When $m = 1$, $k = n$, and $\mathbf{P} = \mathbf{e}^\top$, where $\mathbf{e}$ is a vector of 1s. The strategy is to show that when $m = 1$, $\widetilde{\widehat{\mathbf{B}}}{}^*\mathbf{e}^\top = \mathbf{e}^\top\widetilde{\mathbf{A}}^n$, which will imply that $\tilde{\rho}^2 = 1$ in Equation (11) of the main text. Following this strategy, and using $\widetilde{\widehat{\mathbf{B}}}{}^*$ (the balanced form of the aggregated matrix $\widehat{\mathbf{B}}^*$) yields

$$\begin{aligned}
\widetilde{\widehat{\mathbf{B}}}{}^*\mathbf{e}^\top &= \mathbf{e}^\top\widetilde{\mathbf{A}}^n\mathbf{VWe}(\mathbf{e}^\top\mathbf{VWe})^{-1}\mathbf{e}^\top \\
&= \frac{\mathbf{e}^\top\widetilde{\mathbf{A}}^n(\mathbf{v}\circ\mathbf{w})\mathbf{e}^\top}{\mathbf{v}^\top\mathbf{w}} \\
&= \mathbf{e}^\top\lambda^n(\mathbf{v}\circ\mathbf{w})\mathbf{e}^\top \\
&= \lambda^n(\mathbf{e}^\top)(\mathbf{v}\circ\mathbf{w})\mathbf{e}^\top \\
&= \lambda^n(\mathbf{v}^\top\mathbf{w})\mathbf{e}^\top \\
&= \lambda^n\mathbf{e}^\top \\
&= \mathbf{e}^\top\widetilde{\mathbf{A}}^n,
\end{aligned} \tag{B.1}$$

which demonstrates that, by Equation (11) of the main text, $\tilde{\rho}^2 = 1$. In line 2 of Equation (B.1), we used the identity $\mathbf{e}^\top\mathbf{VWe} = \mathbf{v}^\top\mathbf{w}$ (which is equal to 1), and the identity $\mathbf{VWe} = \mathbf{v}\circ\mathbf{w}$. In line 3, we used the fact that $\mathbf{v}\circ\mathbf{w}$ is an eigenvector of $\widetilde{\mathbf{A}}^n$ associated with dominant eigenvalue $\lambda^n$. In line 5, we used the identity $\mathbf{e}^\top(\mathbf{v} \circ \mathbf{w}) = \mathbf{v}^\top\mathbf{w}$. The final line follows because $\mathbf{e}^\top$ is a left eigenvector of $\widetilde{\mathbf{A}}^{\mathrm{n}}$ associated with eigenvalue $\lambda^n$.

### APPENDIX C Properties of the elasticity-consistent aggregated matrix

#### C.1 The elasticity-consistent aggregated matrix is a Leslie matrix

For $\widehat{\mathbf{B}}^*$ to be a Leslie matrix, the following must be true: (a) it is nonnegative, (b) the subdiagonal entries (survival probabilities) lie between 0 and 1, (c) the entries below the subdiagonal are zero, and (d) the entries above the subdiagonal and below the first row are zero. $\widehat{\mathbf{B}}^*$ is nonnegative because it is the product of nonnegative matrices, so (a) holds.

Recall that $\widehat{\mathbf{B}}^* = \mathbf{P}^*\mathbf{A}\mathbf{Q}$, where $\mathbf{P}^* = \mathbf{P}\mathbf{W}\mathbf{P}^\top(\mathbf{P}\mathbf{V}\mathbf{W}\mathbf{P}^\top)^{-1}\mathbf{P}\mathbf{V}$ and $\mathbf{Q} = \mathbf{W}\mathbf{P}^\top(\mathbf{P}\mathbf{W}\mathbf{P}^\top)^{-1}$. The $m \times n$ matrix $\mathbf{P}^*$ is of the form

$$\mathbf{P}^* = \begin{bmatrix} \mathbf{p}_1^{*\top} & \mathbf{0} & \cdots & \mathbf{0} \\ \mathbf{0} & \mathbf{p}_2^{*\top} & \cdots & \mathbf{0} \\ \vdots & \vdots & \ddots & \vdots \\ \mathbf{0} & \mathbf{0} & \dots & \mathbf{p}_m^{*\top} \end{bmatrix}, \tag{C.1}$$

$$\mathbf{p}_i^{*\top} = [v_{(i-1)k+1} \quad v_{(i-1)k+2} \quad \cdots \quad v_{ik}] \sum_{j=(i-1)k+1}^{ik} w_j \Big/ \sum_{j=(i-1)k+1}^{ik} v_j w_j. \tag{C.2}$$

The matrix $\mathbf{Q}$ is a $n \times m$ matrix of the form has the following form:

$$\mathbf{Q} = \begin{bmatrix} \mathbf{q}_1 & \mathbf{0} & \cdots & \mathbf{0} \\ \mathbf{0} & \mathbf{q}_2 & \cdots & \mathbf{0} \\ \vdots & \vdots & \ddots & \vdots \\ \mathbf{0} & \mathbf{0} & \dots & \mathbf{q}_m \end{bmatrix}, \tag{C.3}$$

where each $k$-vector $\mathbf{q}_i$ is of the form

$$\mathbf{q}_i = [w_{(i-1)k+1} \quad w_{(i-1)k+2} \quad \cdots \quad w_{ik}]^\top / \sum_{j=(i-1)k+1}^{ik} w_j. \tag{C.4}$$

The components of $\mathbf{q}_i$ sum to one, making $\mathbf{Q}$ column stochastic.

We now show that (b) the subdiagonal entries of $\widehat{\mathbf{B}}^*$ lie between zero and one. Consider the entry at row two and column one of the matrix $\widehat{\mathbf{B}}^*$, which we denote $\hat{b}_{21}^*$.

$$
\begin{aligned}
\hat{b}_{21}^{*} &= \mathbf{e}_2^{\top}\widehat{\mathbf{B}}^{*}\mathbf{e}_1 \qquad \text{(C.5)}\\
&= \mathbf{e}_2^{\top}\mathbf{P}^{*}\mathbf{A}^{k}\mathbf{Q}\mathbf{e}_1\\
&= [\mathbf{0} \quad \mathbf{p}_2^{*\top} \quad \mathbf{0} \quad \dots \quad \mathbf{0}]\mathbf{A}^{k}\begin{bmatrix}\mathbf{q}_1\\ \mathbf{0}\\ \mathbf{0}\\ \vdots\\ \mathbf{0}\end{bmatrix}\\
&= (p_{21}^{*}q_{11}P_1P_2\cdots P_k + p_{22}^{*}q_{12}P_2P_3\cdots P_{k+1} + \cdots + p_{2k}^{*}q_{1k}P_kP_{k+1}\cdots P_{2k-1})\\
&= \frac{\sum_{j=1}^{k} v_{k+j}w_{\mathrm{j}}P_{\mathrm{j}}P_2\cdots P_{j+k-1}\sum_{j=k+1}^{2k} w_j}{\sum_{j=1}^{k} w_j\sum_{j=k+1}^{2k} v_jw_j}\\
&= \frac{\lambda^{k}\sum_{j=k+1}^{2k} v_jw_j\sum_{j=k+1}^{2k} w_j}{\sum_{j=1}^{k} w_j\sum_{j=k+1}^{2k} v_jw_j}\\
&= \frac{\lambda^{k}\sum_{j=k+1}^{2k} w_j}{\sum_{j=1}^{k} w_j}\\
&= \frac{\sum_{j=1}^{k} w_{\mathrm{j}}P_{\mathrm{j}}P_2\cdots P_{j+k-1}}{\sum_{j=1}^{k} w_j},
\end{aligned}
$$

where $m$-vector $\mathbf{e}_i$ is a standard basis vector (i.e. it has a one as its $i$th component and zeros elsewhere), $p_{2j}^{*}$ is the $j$th component of $\mathbf{p}_2^{*}$, and $p_{1j}$ is the $j$th component of $\mathbf{q}_1$. Equation (C.5) demonstrates that $\hat{b}_{21}^{*}$ is a convex combination of survival probabilities, which is guaranteed to lie between zero and one. Similarly, $\hat{b}_{32}^{*}, \hat{b}_{43}^{*}, \dots, \hat{b}_{m,m-1}^{*}$ lie between zero and one. The survival probabilities of the elasticity-consistent aggregated matrix are the same as the survival probabilities of the standard aggregated matrix (Hinrichsen, 2023).

Next, we show that (c) the entries of $\widehat{\mathbf{B}}^{*}$ below its subdiagonal are zero. Consider the entry in row 3

$$\hat{b}^*_{31} = \mathbf{e}_3^\top \widehat{\mathbf{B}}^* \mathbf{e}_1 = \mathbf{e}_3^\top \mathbf{P}^* \mathbf{A}^k \mathbf{Q} \mathbf{e}_1 \tag{C.6}$$

$$= [\mathbf{0} \quad \mathbf{0} \quad \mathbf{p}_3^{*\top} \quad \dots \quad \mathbf{0}] \mathbf{A}^k \begin{bmatrix} \mathbf{q}_1 \\ \mathbf{0} \\ \mathbf{0} \\ \vdots \\ \mathbf{0} \end{bmatrix}$$

$$= [\mathbf{0} \quad \mathbf{0} \quad \mathbf{p}_3^{*\top} \quad \dots \quad \mathbf{0}] \begin{bmatrix} * \\ * \\ \mathbf{0} \\ \vdots \\ \mathbf{0} \end{bmatrix}$$

$$= 0.$$

(The $k$-vectors denoted by * are irrelevant to the argument because they are multiplied by zero in the inner product). There are zeros below the first $2k$ components of the $n$-vector

$$\begin{bmatrix} * \\ * \\ \mathbf{0} \\ \vdots \\ \mathbf{0} \end{bmatrix} = \mathbf{A}^k \begin{bmatrix} \mathbf{q}_1 \\ \mathbf{0} \\ \mathbf{0} \\ \vdots \\ \mathbf{0} \end{bmatrix}, \tag{C.7}$$

because multiplying by $\mathbf{A}^k$ advances the first $k$ age classes only as far as the second set of $k$ age classes. Similarly, the remaining entries of $\widehat{\mathbf{B}}^*$ below its subdiagonal are zero.

Lastly, we show that (d) the entries of $\widehat{\mathbf{B}}^*$ that lie above the subdiagonal and below the first row are zero. The entry of $\widehat{\mathbf{B}}^*$in row two and column two is

$$\hat{b}^*_{22} = \mathbf{e}_2^\top \widehat{\mathbf{B}}^* \mathbf{e}_2 = \mathbf{e}_2^\top \mathbf{P}^* \mathbf{A}^k \mathbf{Q} \mathbf{e}_2 \tag{C.8}$$

$$= [\mathbf{0} \quad \mathbf{p}_2^{*\top} \quad \mathbf{0} \quad \dots \quad \mathbf{0}] \mathbf{A}^k \begin{bmatrix} \mathbf{0} \\ \mathbf{q}_2 \\ \mathbf{0} \\ \vdots \\ \mathbf{0} \end{bmatrix}$$

$$= [\mathbf{0} \quad \mathbf{p}_2^{*\top} \quad \mathbf{0} \quad \dots \quad \mathbf{0}] \begin{bmatrix} * \\ \mathbf{0} \\ * \\ \vdots \\ \mathbf{0} \end{bmatrix}$$

$$= 0.$$

There are *k* zeroes immediately after the first *k* components of

$$\begin{bmatrix} * \\ \mathbf{0} \\ * \\ \vdots \\ \mathbf{0} \end{bmatrix} = \mathbf{A}^k \begin{bmatrix} \mathbf{0} \\ \mathbf{q}_2 \\ \mathbf{0} \\ \vdots \\ \mathbf{0} \end{bmatrix}, \tag{C.9}$$

because multiplication by $\mathbf{A}^k$ advances individuals in the second set of *k* age classes to the third set of *k* age classes, and after *k* time steps, the offspring of these individuals are too young to advance to the second set of *k* age classes. Similarly, the remaining entries of $\widehat{\mathbf{B}}^*$ above the subdiagonal and below the first row are zero. Thus, properties (a)–(d) hold, demonstrating that $\widehat{\mathbf{B}}^*$ is a Leslie matrix.

### C.2 The aggregated matrix is irreducible whenever A is irreducible

A Leslie matrix is irreducible precisely when the survival probabilities are all positive and fertility rate of the oldest age class is positive (Freese & Johnson, 1974). Therefore, to demonstrate that $\widehat{\mathbf{B}}^*$ is irreducible if **A** is irreducible, we need only show that $\hat{b}^*_{21}, \hat{b}^*_{32}, \ldots, \hat{b}^*_{m,m-1}$ (i.e. the survival probabilities in matrix $\widehat{\mathbf{B}}^*$ ) are all positive and $\hat{b}^*_{1,m} > 0$ if the age-specific survival probabilities, $P_1, P_2, \ldots, P_{n-1}$ are all positive and $F_n > 0$.

Assume that **A** is irreducible so that the survival probabilities in the matrix **A** are positive and $F_n > 0$. We first show that the survival probabilities in matrix $\widehat{\mathbf{B}}^*$ are also positive. Using Equation (C.5):

$$\hat{b}^*_{21} = \frac{\sum_{j=1}^{k} w_{\mathrm{j}} P_{\mathrm{j}} P_2 \cdots P_{j+k-1}}{\sum_{j=1}^{k} w_j} \tag{C.10}$$

Note that $w_{\mathrm{j}}$ is positive since it is a component of **w**, the stable age distribution, which is guaranteed to be positive. Furthermore, $P_j, P_2, \ldots, P_{j+k-1}$ are positive. Therefore, $\hat{b}^*_{21} > 0$. Similar arguments show that the rest of the survival probabilities in $\widehat{\mathbf{B}}^*$ are positive.

Next, we show that $\hat{b}^*_{1m}$ is positive. By the Perron-Frobenius theorem, if **A** is nonnegative and irreducible, its reproductive values and stable age distribution are positive, so that $\mathbf{v} > 0$ and $\mathbf{w} > 0$, and therefore, $\mathbf{p}^*_i > 0$ and $\mathbf{q}_i > 0$ for $j = 1,2, \ldots m$.

$$\begin{aligned}
\hat{b}^*_{1m} &= \mathbf{e}_1^\top \widehat{\mathbf{B}}^* \mathbf{e}_m \\
&= \mathbf{e}_1^\top \mathbf{P}^* \mathbf{A}^k \mathbf{Q} \mathbf{e}_m \\
&= [\mathbf{p}_1^{*\top} \quad \mathbf{0} \quad \mathbf{0} \quad \dots \quad \mathbf{0}] \mathbf{A}^k \begin{bmatrix} \mathbf{0} \\ \mathbf{0} \\ \mathbf{0} \\ \vdots \\ \mathbf{q}_m \end{bmatrix} \\
&\geq [\mathbf{p}_1^{*'\top} \quad \mathbf{0} \quad \mathbf{0} \quad \dots \quad \mathbf{0}] \begin{bmatrix} 0 & 0 & \cdots & 0 & F_n \\ P_1 & 0 & \cdots & 0 & 0 \\ 0 & P_2 & \cdots & 0 & 0 \\ \vdots & \vdots & \ddots & \vdots & \vdots \\ 0 & 0 & \dots & P_{n-1} & 0 \end{bmatrix}^k \begin{bmatrix} \mathbf{0} \\ \mathbf{0} \\ \mathbf{0} \\ \vdots \\ \mathbf{q}'_m \end{bmatrix} \\
&= p^*_{1,k} q_{m,k} F_n P_1 P_2 \dots P_{k-1},
\end{aligned} \tag{C.11}$$

where $\mathbf{p}_1^{*'\top} = [0 \quad 0 \quad \dots \quad p^*_{1,k}]$ and $\mathbf{q}'_m = [0 \quad 0 \quad \dots \quad q_{m,k}]^\top$. Line 4 of Equation (C.11) was produced by replacing **A** with a Leslie matrix $\mathbf{A}'$ that has fertility rates that are zero except for the last age class, which has the same fertility rate as **A**, namely, $F_n$. We also replaced $\mathbf{p}_1^{*\top}$ with $\mathbf{p}_1^{*'\top}$ and $\mathbf{q}_m$ with $\mathbf{q}'_m$. Because each replacement is less than the matrix or vector that it replaces, the product in line 4 of Equation (C.11) is less than or equal to $\hat{b}^*_{1m}$. Since **A** was assumed irreducible, $p^*_{1,k} > 0$, $q_{m,k} > 0, F_n > 0$, and $P_1 P_2 \dots P_{k-1} > 0$, and therefore, $p^*_{1,k} q_{m,k} F_n P_1 P_2 \dots P_{k-1} > 0$, and by Equation (C.11), $\hat{b}^*_{1m} > 0$. Since its survival probabilities are positive and the fertility rate of its oldest age class is positive, $\widehat{\mathbf{B}}^*$ is irreducible (Freese & Johnson, 1974).

#### C.3 The elasticity-consistent aggregated matrix is primitive whenever A is primitive

We next show that if $\mathbf{A}$ is primitive, then $\widehat{\mathbf{B}}^*$is primitive too. A nonnegative matrix is primitive if raising it to some integer power greater than zero produces a matrix whose entries are all positive (Keyfitz & Caswell, 2005). Assume that $\mathbf{A}$ is primitive. Then $\mathbf{A}^k$ is also primitive (Horn & Johnson, 2013). Since $\mathbf{A}^k$ is primitive, then $\mathbf{A}^{ks} > 0$ for some integer $s \geq 1$. To demonstrate that $\widehat{\mathbf{B}}^*$ is also primitive, the following inequality is useful:

$$\mathbf{QP}^* = \begin{bmatrix} \mathbf{q_1} & \mathbf{0} & \cdots & \mathbf{0} \\ \mathbf{0} & \mathbf{q_2} & \cdots & \mathbf{0} \\ \vdots & \vdots & \ddots & \vdots \\ \mathbf{0} & \mathbf{0} & \ldots & \mathbf{q}_m \end{bmatrix} \begin{bmatrix} \mathbf{p}_1^{*\top} & \mathbf{0} & \cdots & \mathbf{0} \\ \mathbf{0} & \mathbf{p}_2^{*\top} & \cdots & \mathbf{0} \\ \vdots & \vdots & \ddots & \vdots \\ \mathbf{0} & \mathbf{0} & \ldots & \mathbf{p}_m^{*\top} \end{bmatrix} \tag{C.12}$$

$$= \begin{bmatrix} \mathbf{q}_1\mathbf{p}_1^{*\top} & \mathbf{0} & \cdots & \mathbf{0} \\ \mathbf{0} & \mathbf{q}_2\mathbf{p}_1^{*\top} & \cdots & \mathbf{0} \\ \vdots & \vdots & \ddots & \vdots \\ \mathbf{0} & \mathbf{0} & \ldots & \mathbf{q}_m\mathbf{p}_1^{*\top} \end{bmatrix}$$

$$\geq \gamma\mathbf{I},$$

where $\gamma$ is a positive scalar equal to the smallest entry in the set of matrices $\mathbf{q}_1\mathbf{p}_1^{*\top}, \mathbf{q}_2\mathbf{p}_1^{*\top}, \ldots, \mathbf{q}_m\mathbf{p}_m^{*\top}$ . Since $\mathbf{A}$ is primitive, $\mathbf{v} > 0$ and $\mathbf{w} > 0$, and therefore, $\mathbf{p}_i^* > 0$ and $\mathbf{q}_i > 0$ for $j = 1,2, \ldots m$, and $\gamma > 0$.

Writing $\widehat{\mathbf{B}}^{*s}$ as a product of $s$ copies of $\widehat{\mathbf{B}}^* = \mathbf{P}^*\mathbf{A}^k\mathbf{Q}$ gives

$$\begin{aligned}\widehat{\mathbf{B}}^{*s} &= (\mathbf{P}^*\mathbf{A}^k\mathbf{Q})^s \\ &= (\mathbf{P}^*\mathbf{A}^k\mathbf{Q})(\mathbf{P}^*\mathbf{A}^k\mathbf{Q})\cdots(\mathbf{P}^*\mathbf{A}^k\mathbf{Q}) \\ &= \mathbf{P}^*\mathbf{A}^k(\mathbf{Q}\,\mathbf{P}^*)\mathbf{A}^k(\mathbf{Q}\,\mathbf{P}^*)\cdots\mathbf{A}^k(\mathbf{Q}\,\mathbf{P}^*)\mathbf{A}^k\mathbf{Q} \\ &\geq \mathbf{P}^*\mathbf{A}^k(\gamma\mathbf{I})\mathbf{A}^k(\gamma\mathbf{I})\cdots\mathbf{A}^k(\gamma\mathbf{I})\mathbf{A}^k\mathbf{Q} \\ &= \gamma^{s-1}\mathbf{P}^*\mathbf{A}^{ks}\mathbf{Q}.\end{aligned} \tag{C.13}$$

The inequality in line 4 of Equation (C.13) holds because we replaced each $\mathbf{Q}\,\mathbf{P}^*$ with $\gamma\mathbf{I}$, and by Equation (C.12), $\mathbf{Q}\,\mathbf{P}^* \geq \gamma\mathbf{I}$. The entries of $\mathbf{P}^*\mathbf{A}^{ks}\mathbf{Q}$ are positive because each entry is a linear combination of the entries of a $k \times k$ positive submatrix of $\mathbf{A}^{ks}$ with positive weights from $\mathbf{P}^*$ and $\mathbf{Q}$:

$$\begin{aligned}&\mathbf{P}^*\mathbf{A}^{ks}\mathbf{Q} \\ &= \begin{bmatrix}\mathbf{p}_1^{*\top} & \mathbf{0} & \cdots & \mathbf{0} \\ \mathbf{0} & \mathbf{p}_2^{*\top} & \cdots & \mathbf{0} \\ \vdots & \vdots & \ddots & \vdots \\ \mathbf{0} & \mathbf{0} & \dots & \mathbf{p}_m^{*\top}\end{bmatrix}\begin{bmatrix}(\mathbf{A}^{ks})_{11} & (\mathbf{A}^{ks})_{12} & \cdots & (\mathbf{A}^{ks})_{1m} \\ (\mathbf{A}^{ks})_{21} & (\mathbf{A}^{ks})_{22} & \cdots & (\mathbf{A}^{ks})_{2m} \\ \vdots & \vdots & \ddots & \vdots \\ (\mathbf{A}^{ks})_{m1} & (\mathbf{A}^{ks})_{m2} & \dots & (\mathbf{A}^{ks})_{mm}\end{bmatrix}\begin{bmatrix}\mathbf{q}_1 & \mathbf{0} & \cdots & \mathbf{0} \\ \mathbf{0} & \mathbf{q}_2 & \cdots & \mathbf{0} \\ \vdots & \vdots & \ddots & \vdots \\ \mathbf{0} & \mathbf{0} & \dots & \mathbf{q}_m\end{bmatrix} \\ &= \begin{bmatrix}\mathbf{p}_1^{*\top}(\mathbf{A}^{ks})_{11}\mathbf{q}_1 & \mathbf{p}_1^{*\top}(\mathbf{A}^{ks})_{12}\mathbf{q}_2 & \cdots & \mathbf{p}_1^{*\top}(\mathbf{A}^{ks})_{1m}\mathbf{q}_m \\ \mathbf{p}_2^{*\top}(\mathbf{A}^{ks})_{21}\mathbf{q}_1 & \mathbf{p}_2^{*\top}(\mathbf{A}^{ks})_{22}\mathbf{q}_2 & \cdots & \mathbf{p}_2^{*\top}(\mathbf{A}^{ks})_{2m}\mathbf{q}_m \\ \vdots & \vdots & \ddots & \vdots \\ \mathbf{p}_m^{*\top}(\mathbf{A}^{ks})_{m1}\mathbf{q}_1 & \mathbf{p}_m^{*\top}(\mathbf{A}^{ks})_{m2}\mathbf{q}_2 & \dots & \mathbf{p}_m^{*\top}(\mathbf{A}^{ks})_{mm}\mathbf{q}_m\end{bmatrix},\end{aligned} \tag{C.14}$$

where $(\mathbf{A}^{ks})_{ij}$ is a $k \times k$ block (submatrix) of $\mathbf{A}^{ks}$. Therefore $\gamma^{s-1}\mathbf{P}^*\mathbf{A}^{ks}\mathbf{Q} > 0$, and by Equation (C.13), $\widehat{\boldsymbol{B}}^{*s} > 0$, so $\widehat{\boldsymbol{B}}^*$ is primitive.

### APPENDIX D Aggregating with a disaggregation step

#### D.1 Disaggregated matrix

When $m$ does not evenly divide $n$, it is still possible to aggregate the matrix **A** into a smaller $m \times m$ matrix. To proceed, we first disaggregate the original Leslie matrix **A** into a larger Leslie matrix **C** that is consistent with the original Leslie matrix. We construct the disaggregated Leslie matrix **C** so that its dimensionality is a multiple of $m$, then apply the usual aggregator (the elasticity-consistent aggregator in this case) to obtain the aggregated matrix $\widehat{\mathbf{B}}_d^*$**.** **(**The subscript $d$ is used to denote an aggregated matrix derived from a disaggregation step.)

Such a Leslie matrix $\mathbf{C}$ is easily constructed and always exists (Hinrichsen, 2023). To construct the disaggregated Leslie matrix $\mathbf{C}$, subdivide each age class of **A** into $m$ sub classes to obtain

$$\mathbf{C} = \begin{bmatrix} c_{11} & c_{12} & c_{13} & \cdots & c_{1,mn-1} & c_{1,mn} \\ c_{21} & 0 & 0 & \cdots & 0 & 0 \\ 0 & c_{32} & 0 & \cdots & 0 & 0 \\ 0 & 0 & c_{43} & \cdots & 0 & 0 \\ \vdots & \vdots & \vdots & \ddots & \vdots & \vdots \\ 0 & 0 & 0 & \dots & c_{mn,mn-1} & 0 \end{bmatrix}, \tag{D.1}$$

where $c_{1j} = 0$ if $j$ is not a multiple of $m$; $c_{1j} = F_l$ if $j = lm$, where $l$ is an integer; $c_{i+1,i} = 1$ if $i$ is not a multiple of $m$; and $c_{i+1,i} = P_l$ if $i = lm$, where $l$ is an integer. Raising $\mathbf{C}$ to the $m$th power yields,

$$\mathbf{C}^m = \mathbf{A} \otimes \mathbf{I}_m = \begin{bmatrix} F_1\mathbf{I}_m & F_2\mathbf{I}_m & F_3\mathbf{I}_m & \cdots & F_{n-1}\mathbf{I}_m & F_n\mathbf{I}_m \\ P_1\mathbf{I}_m & 0 & 0 & \cdots & 0 & 0 \\ 0 & P_2\mathbf{I}_m & 0 & \cdots & 0 & 0 \\ 0 & 0 & P_3\mathbf{I}_m & \cdots & 0 & 0 \\ \vdots & \vdots & \vdots & \ddots & \vdots & \vdots \\ 0 & 0 & 0 & \dots & P_{n-1}\mathbf{I}_m & 0 \end{bmatrix}, \tag{D.2}$$

where $\otimes$ denotes the Kronecker product (Horn & Johnson, 1991). Equation (D.2) shows that $\mathbf{C}^m$ contains exactly $m$ copies of the original Leslie matrix $\mathbf{A}$. Therefore, the eigenvalues of $\mathbf{A}$ and $\mathbf{C}^m$ are the same, which follows because the eigenvalues of $\mathbf{C}^m = \mathbf{A} \otimes \mathbf{I}_m$ are products of the eigenvalues of $\mathbf{A}$ and the eigenvalues of $\mathbf{I}_m$ (Horn & Johnson, 1991). Because the eigenvalues of $\mathbf{I}_m$ are all 1, the eigenvalues of $\mathbf{A}$ and $\mathbf{C}^m$ are the same, and the algebraic multiplicity of each eigenvalue of $\mathbf{C}^m$ equals $m$ times its algebraic multiplicity as an eigenvalue in $\mathbf{A}$.

The original matrix $\mathbf{A}$ is a perfect aggregation of the matrix $\mathbf{C}^m$ associated with the $n \times mn$ partitioning matrix

$$\mathbf{P}_1 = \begin{bmatrix} \mathbf{e}^\top & \mathbf{0} & \cdots & \mathbf{0} \\ \mathbf{0} & \mathbf{e}^\top & \cdots & \mathbf{0} \\ \vdots & \vdots & \ddots & \vdots \\ \mathbf{0} & \mathbf{0} & \dots & \mathbf{e}^\top \end{bmatrix}, \tag{D.3}$$

where $\boldsymbol{e}$ is an $m$-vector of ones. Perfect aggregation holds because

$$\mathbf{P}_1\mathbf{C}^m = \mathbf{P_1}(\mathbf{A} \otimes \mathbf{I}_m) \qquad \text{(D.4)}$$
$$= \mathbf{A} \otimes \boldsymbol{e}^\top$$
$$= \mathbf{A}\mathbf{P}_1.$$

Since $\mathbf{A}$ is a perfect aggregation of the matrix $\mathbf{C}^m$, the dynamics of the original system governed by $\mathbf{A}$ are exactly represented with the larger disaggregated system governed by $\mathbf{C}^m$. That is, if we begin with an individual in age class $i$ of $\mathbf{A}$, and determine that individual's contribution to age class $j$ of $\mathbf{A}$ at time $t$, this is equivalent to beginning with an individual in set $i$ of $m$ age classes of $\mathbf{C}$, determining its contribution to set $j$ of $m$ age classes of $\mathbf{C}$ at time $t$. The exponent of $m$ is used for $\mathbf{C}$ because the projection interval of $\mathbf{A}$ is $m$ times the projection interval of $\mathbf{C}$.

The elasticity-consistent aggregated matrix is

$$\widehat{\mathbf{B}}_d^* = \mathbf{P}_2^*\mathbf{C}^n\mathbf{Q}_2, \qquad \text{(D.5)}$$

where the $m \times nm$ partitioning matrix $\mathbf{P}_2^*$ and the $nm \times m$ matrix $\mathbf{Q}_2$ are the analogues of $\mathbf{P}^*$ and $\mathbf{Q}$, respectively, in Equation (10) of the main text.

### D.2 Properties of elasticity-consistent aggregated matrix constructed using a disaggregation step

The elasticity-consistent aggregated matrix derived using the disaggregation step, $\widehat{\mathbf{B}}_d^*$, inherits the following properties from the original Leslie matrix **A**: it is an $m \times m$ Leslie matrix, it is irreducible whenever **A** is irreducible, and it is primitive whenever **A** is primitive. Furthermore, the stable growth rate, stable age distribution, and reproductive values of the aggregated system are consistent with those of the original Leslie model. These are the same properties enjoyed by $\widehat{\mathbf{B}}^*$, derived when $m$ divides $n$ evenly.

Since $\widehat{\mathbf{B}}_d^*$ is the elasticity-consistent aggregated matrix formed from the matrix Leslie matrix **C**, $\widehat{\mathbf{B}}_d^*$ is a Leslie matrix and $\widehat{\mathbf{B}}_d^*$ is irreducible whenever **C** is irreducible (see Appendix D). Therefore, to demonstrate these properties for $\widehat{\mathbf{B}}_d^*$, we need only show that **C** is a Leslie matrix and is irreducible whenever **A** is irreducible. The disaggregated matrix **C** is indeed a Leslie matrix by the way it was defined. **C** is also irreducible whenever **A** is irreducible because all the survival probabilities in **C** are positive and $c_{1,mn} = F_n$. If **A** is irreducible, $F_n > 0$, and therefore, $c_{1,mn} > 0$. Since a Leslie matrix is irreducible if and only if its survival probabilities are positive and the fertility rate of its oldest age class is positive (Freese & Johnson, 1974), **C** is irreducible.

We next show that $\widehat{\mathbf{B}}_d^*$ is primitive whenever **A** is primitive. The argument is similar to that given in Appendix D. A nonnegative matrix is primitive if raising it to some integer power greater than zero yields a matrix with entries that are all positive (Keyfitz & Caswell, 2005).

Assume that $\mathbf{A}$ is primitive, then $\mathbf{A}^n$ is also primitive (Horn & Johnson, 2013). Since $\mathbf{A}^n$ is primitive, then $\mathbf{A}^{ns_2} > 0$ for some $s_2 \geq 1$.

As in Equation (C.12) there exists a positive constant $\gamma_2$ such that

$$\mathbf{Q}_2\mathbf{P}_2^* \geq \gamma_2\mathbf{I}_{mn}, \tag{D.6}$$

where $\mathbf{I}_{mn}$ is an $mn \times mn$ identity matrix.

$$\begin{aligned}
\widehat{\mathbf{B}}_d^{*ms_2} &= (\mathbf{P}_2^*\mathbf{C}^n\mathbf{Q}_2)^{ms_2} \\
&= (\mathbf{P}_2^*\mathbf{C}^n\mathbf{Q}_2)(\mathbf{P}_2^*\mathbf{C}^n\mathbf{Q}_2)\cdots(\mathbf{P}_2^*\mathbf{C}^n\mathbf{Q}_2) \\
&= \mathbf{P}_2^*\mathbf{C}^n(\mathbf{Q}_2\,\mathbf{P}_2^*)\mathbf{C}^n(\mathbf{Q}_2\,\mathbf{P}_2^*)\cdots\mathbf{C}^n(\mathbf{Q}_2\,\mathbf{P}_2^*)\mathbf{C}^n\mathbf{Q}_2 \\
&\geq \gamma^{ms_2-1}\mathbf{P}_2^*\mathbf{C}^{mns_2}\mathbf{Q}_2 \\
&= \gamma^{ms_2-1}\mathbf{P}_2^*(\mathbf{A}\otimes\mathbf{I}_m)^{ns_2}\mathbf{Q}_2 \\
&= \gamma^{ms_2-1}\mathbf{P}_2^*(\mathbf{A}^{ns_2}\otimes\mathbf{I}_m)\mathbf{Q}_2.
\end{aligned} \tag{D.7}$$

Line 5 of Equation (D.7) follows because $\mathbf{C}^m = \mathbf{A}\otimes\mathbf{I}_m$. The last line of Equation (D.7) follows from the properties of the Kronecker product. Since $\mathbf{A}^{ns_2} > \mathbf{0}$ (by assumption), $\mathbf{A}^{ns_2}\otimes\mathbf{I}_m$ is a block matrix, where each block is an $m \times m$ diagonal matrix with positive entries on its diagonal. Write out the product of $\mathbf{P}_2^*(\mathbf{A}^{ns_2}\otimes\mathbf{I}_m)\mathbf{Q}_2$ to derive an equation like Equation (C.14). Each $n \times n$ block of $\mathbf{A}^{ns_2}\otimes\mathbf{I}_m$ must contain a positive entry because $m < n$ and the largest square block of all zeroes in $\mathbf{A}^{nq_2} \otimes \mathbf{I}_m$ is of size $\lfloor m/2 \rfloor \times \lfloor m/2 \rfloor$, where $\lfloor \cdot \rfloor$ is the floor function. Therefore, as in Equation (C.14), multiplying $\mathbf{A}^{ns_2} \otimes \mathbf{I}_m$ on the left by $\mathbf{P}_2^*$ and on the right by $\mathbf{Q}_2$ reduces each of these $n \times n$ blocks to a positive number. Therefore, $\mathbf{P}_2^*(\mathbf{A}^{ns_2} \otimes \mathbf{I}_m)\mathbf{Q}_2 > \mathbf{0}$, and by Equation (D.7), $\widehat{\mathbf{B}}_d^{*ms_2} > \mathbf{0}$, and $\widehat{\mathbf{B}}_d^{*m}$ is primitive.

**D.3. Consistency of asymptotic dynamics**

As in the case where $m$ divides $n$ evenly, the elasticity-consistent aggregation matrix $\widehat{\mathbf{B}}_d^*$ has stable growth rate, stable age distribution, and reproductive values that are consistent with the original Leslie matrix, $\mathbf{A}$. We make use of the principle that if $\widehat{\mathbf{B}}_d^*$ is consistent with $\mathbf{C}$ and $\mathbf{A}$ is consistent with $\mathbf{C}$, then $\widehat{\mathbf{B}}_d^*$ is consistent with $\mathbf{A}$ (transitivity). To show consistency of asymptotic dynamics, I use the same arguments in Appendix A but applied to $\mathbf{C}^n$ instead of $\mathbf{A}^k$. We first demonstrate consistency of the dominant eigenvalue and stable age distribution. Using Equation (A.7) and replacing $\widehat{\mathbf{B}}^*$, $\mathbf{A}^k$, $\mathbf{P}$, and $\mathbf{w}$ with $\widehat{\mathbf{B}}_d^*$, $\mathbf{C}^n$, $\mathbf{P}_2$, and $\mathbf{w}_2$, respectively, yields

$$\widehat{\mathbf{B}}_d^*\mathbf{P}_2\mathbf{w}_2 = \lambda(\mathbf{C}^n)\mathbf{P}_2\mathbf{w}_2. \tag{D.8}$$

Since $\widehat{\mathbf{B}}_d^*$ is a nonnegative Leslie matrix, $\mathbf{P}_2\mathbf{w}_2$ must be a right eigenvector of $\widehat{\mathbf{B}}_d^*$ corresponding to the dominant eigenvalue

$$\lambda(\widehat{\mathbf{B}}_d^*) = \lambda(\mathbf{C}^n) \tag{D.9}$$
$$= \lambda(\mathbf{C}^m)^{\frac{n}{m}}$$
$$= \lambda(\mathbf{A})^k.$$

The last line of Equation (D.9) follows because the eigenvalues of $\mathbf{A}$ and $\mathbf{C}^m$ are the same (but with different algebraic multiplicities). Equation (D.9) shows that the dominant eigenvalues of $\mathbf{A}$ and $\widehat{\mathbf{B}}_d^*$ are consistent. The stable age distributions of $\mathbf{A}$ and $\mathbf{C}$ are also consistent because the stable age distribution of $\mathbf{A}$ is $\mathbf{P}_1\mathbf{w}_2$.

To show consistency of the reproductive values of $\widehat{\mathbf{B}}_d^*$ and $\mathbf{C}$, we use Equation (A.8), but applied to $\mathbf{C}^n$ instead of $\boldsymbol{A^k}$ to obtain

$$\mathbf{v}_{\widehat{\mathbf{B}}_d^*} = (\mathbf{P}_2\mathbf{W}_2\mathbf{P}_2^\top)^{-\mathbf{1}}\mathbf{P}_2\mathbf{W}_2\mathbf{v}_2, \tag{D.10}$$

where $\mathbf{v}_{\widehat{\mathbf{B}}_d^*}$ is the vector of reproductive values of $\widehat{\mathbf{B}}_d^*$, $\mathbf{v}_2$ is the vector of reproductive values of $\mathbf{C}$, and $\mathbf{W}_2 = \mathrm{diag}(\mathbf{w}_2)$. With these substitutions, by Equation (A.8), $\mathbf{v}_{\widehat{\mathbf{B}}_d^*}$ is consistent with $\mathbf{v}_2$. The reproductive values of $\mathbf{A}$, are given by

$$\mathbf{v} = (\mathbf{P}_1\mathbf{W}_2\mathbf{P}_1^\top)^{-\mathbf{1}}\mathbf{P}_1\mathbf{W}_2\mathbf{v}_2\ . \tag{D.11}$$

The reproductive values are consistent with those of $\mathbf{C}$ because $\mathbf{A}$ is a perfect aggregation $\mathbf{C}$ associated with $\mathbf{P}_1$.

To summarize, the stable growth rate, stable age distribution, and reproductive values of $\widehat{\mathbf{B}}_d^*$ and $\mathbf{A}$ are consistent with those of $\mathbf{C}$. Therefore, by transitivity, they are also consistent between $\widehat{\mathbf{B}}_d^*$ and $\mathbf{A}$.

**D.4 Example of aggregation using disaggregation step**

Let $\mathbf{A}$ be the $3 \times 3$ $(n = 3)$ matrix

$$\mathbf{A} = \begin{bmatrix} 0 & 1 & 5 \\ .3 & 0 & 0 \\ 0 & .5 & 0 \end{bmatrix}, \tag{D.12}$$

and the goal is to aggregate $\mathbf{A}$ into a $2 \times 2$ $(m = 2)$ matrix. Since $m$ does not divide $n$ evenly, first disaggregate $\mathbf{A}$ into a larger $6 \times 6$ Leslie matrix $\mathbf{C}$ using Equation (D.1):

$$\mathbf{C} = \begin{bmatrix} 0 & 0 & 0 & 1 & 0 & 5 \\ 1 & 0 & 0 & 0 & 0 & 0 \\ 0 & .3 & 0 & 0 & 0 & 0 \\ 0 & 0 & 1 & 0 & 0 & 0 \\ 0 & 0 & 0 & .5 & 0 & 0 \\ 0 & 0 & 0 & 0 & 1 & 0 \end{bmatrix}. \tag{D.13}$$

The $2 \times 2$ elasticity-consistent aggregated matrix is

$$\widehat{\mathbf{B}}_d^* = \mathbf{P}_2^* \mathbf{C}^3 \mathbf{Q}_2 \tag{D.14}$$

$$\cong \begin{bmatrix} 0.20 & 3.26 \\ 0.26 & 0 \end{bmatrix}.$$

Calculating growth rates gives $\lambda(\widehat{\mathbf{B}}_d^*)^{2/3} = \lambda(\mathbf{A}) \cong 1.02$. The exponent of 2/3 is used in the formula because the projection interval of $\widehat{\mathbf{B}}_d^*$ is $k = 3/2$ times the projection interval of $\mathbf{A}$.

### APPENDIX E Transforming Lefkovitch matrices to Leslie matrices

We transformed the original Lefkovitch matrices for soft-shell clam and earthworm into Leslie matrices of dimensionality 11 and 7, respectively. Each original $s \times s$ Lefkovitch matrix, $\mathbf{L} = (l_{\alpha\beta})$, had the form of a Leslie matrix, except that its entry in the southeast corner ($l_{s,s}$) was positive instead of zero. We constructed the $n \times n$ Leslie matrices with $n > s$ as follows:

$$F_i = \begin{cases} l_{1,i} & \text{if } i \leq s \\ l_{1,s} & \text{if } i > s \end{cases} \tag{E.1}$$

and

$$P_j = \begin{cases} l_{j+1,j} & \text{if } j < s \\ l_{s,s} & \text{if } j \geq s \end{cases} \tag{E.2}$$

where $F_i$ $(1 \leq i \leq n)$ were the fertility rates and $P_j$ $(1 \leq j \leq n-1)$ were the survival probabilities in Equation (1) of the main text. We chose the dimensionality of the Leslie matrix, $n$, as the smallest dimensionality such that the stable growth rate of the Leslie matrix approximated that of the original Lefkovitch matrix with relative error less than 0.1%.

### References (Appendices A-E)

Freese, R., & Johnson, C. R. (1974). Comments on the discrete matrix model of population dynamics. *Journal of Research of the National Bureau of Standards, Section B: Mathematical Sciences*, 78B(2), 73–78. https://doi.org/10.6028/jres.078b.012

Golub, G. H., & Van Loan, C. F. (2013). *Matrix computations* (4th ed.). The Johns Hopkins University Press.

Hinrichsen, R. A. (2023). Aggregation of Leslie matrix models with application to ten diverse animal species. *Population Ecology*, 65(3), 146–166. https://doi.org/10.1002/1438-390X.12149

Hinrichsen, R. A. (2025). Scale-invariant interstage flow matrices: a comment on Yokomizo et al. (2024). *Journal of Ecology*, 113(6), 1451–1457.

Horn, R. A., & Johnson, C. R. (1991). *Topics in matrix analysis.* Cambridge University Press. https://doi.org/10.1017/CBO9780511840371

Horn, R. A., & Johnson, C. R. (2013). *Matrix analysis* (2nd ed.). Cambridge University Press. https://doi.org/10.1017/9781139020411

Keyfitz, N., & Caswell, H. (2005). *Applied mathematical demography* (3rd ed.). Springer. https://doi.org/10.1007/b139042

Yokomizo, H., Fukaya, K., Lambrinos, J. G., & Takada, T. (2025). The scale-variant interstage flow makes biological insights possible: A response to Hinrichsen. *Journal of Ecology*, 113(6), 1458–1460. https://doi.org/10.1111/1365-2745.70049

### APPENDIX F Supplemental figures and tables

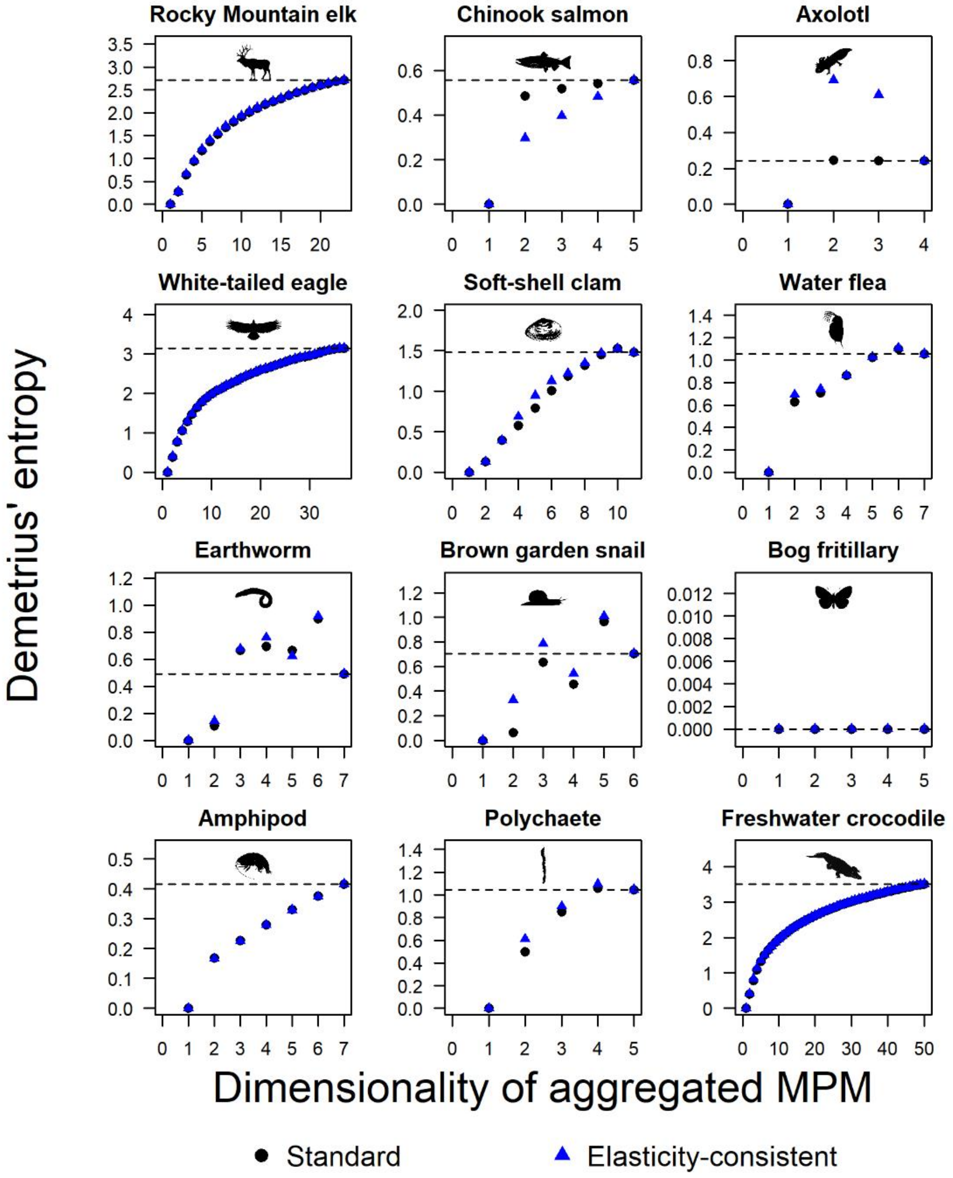


Figure S1: Demetrius' entropy tends to be underestimated with aggregation, but with exceptions (axolotl, earthworm, and bog fritillary). Illustrated is Demetrius' entropy estimated from standard and elasticity-consistent aggregators across 12 animal species, evaluated against the dimensionality of the aggregated matrix population model (MPM), $m$. The dashed horizontal line represents the actual Demetrius' entropy, derived from the original (unaggregated) MPM. Silhouettes are identical to those in Figure 3 of the main text; credits are given in the Figure 3 caption.

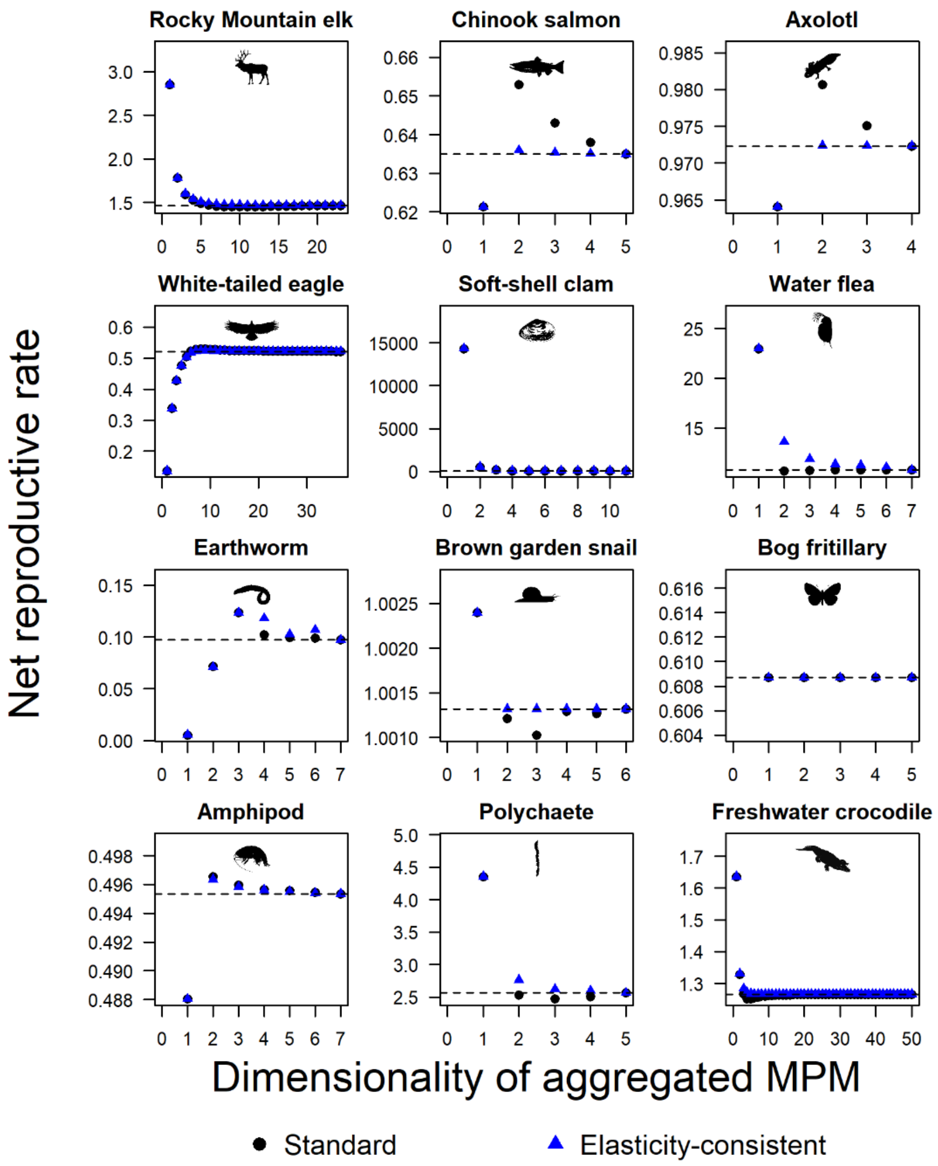


Figure S2: Although stable growth rate is reproduced without error when aggregating, its lifetime reproductive counterpart, net reproductive rate is not (except for bog fritillary which has an associated imprimitive Leslie matrix). Illustrated is the net reproductive rate estimated from standard and elasticity-consistent aggregators across 12 animal species, evaluated against the dimensionality of the aggregated matrix population model (MPM), $m$. The dashed horizontal line represents the actual net reproductive rate, derived from the original (unaggregated) MPM. Silhouettes are identical to those in Figure 3 of the main text; credits are given in the Figure 3 caption.

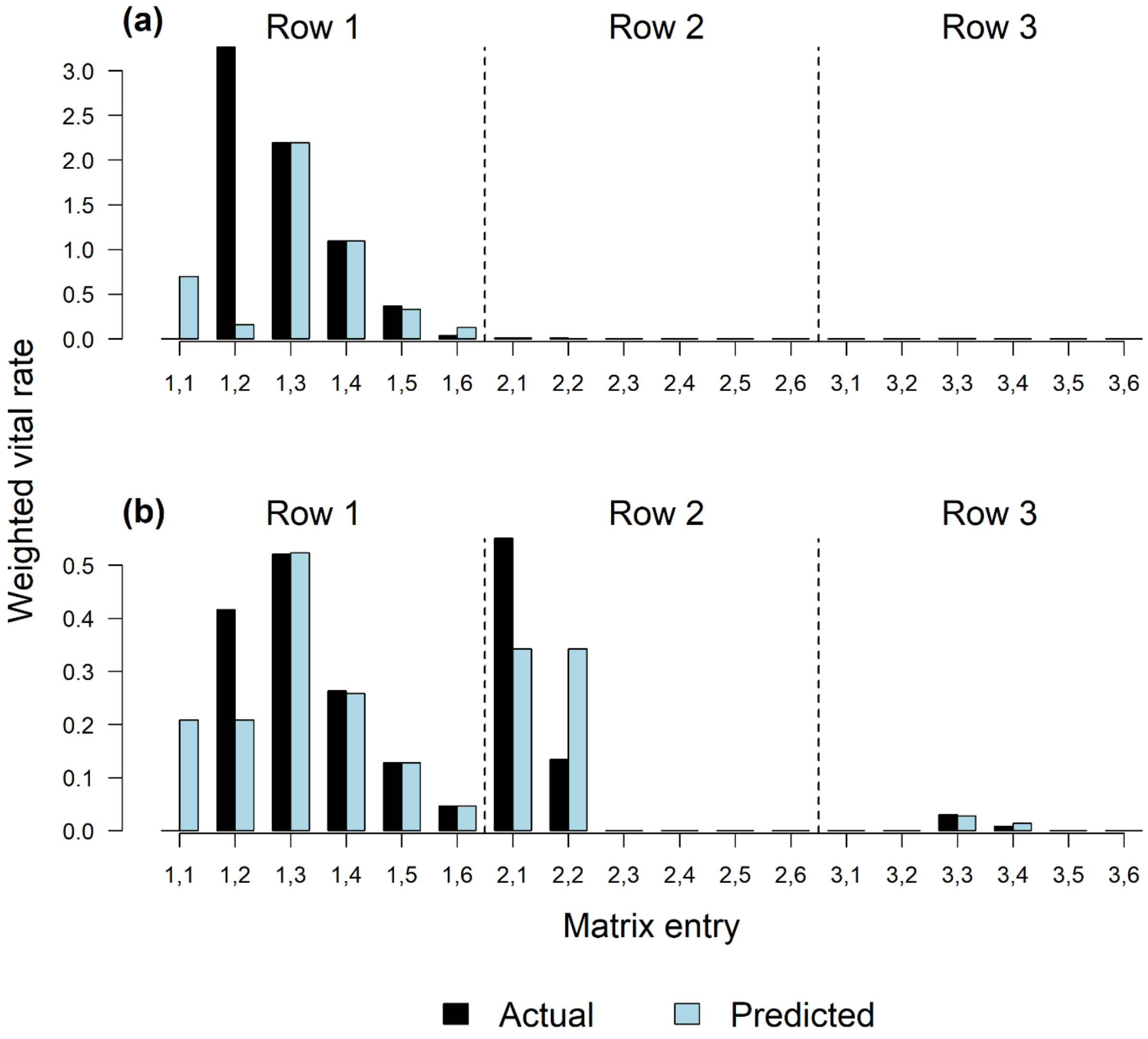


Figure S3: Balancing by reproductive values produces higher effectiveness of aggregation because balanced matrix entries tend to be less variable and therefore easier to fit with a lower dimensional matrix. Illustrated are predicted values compared with actual values for (a) standard aggregation and (b) elasticity-consistent aggregation of a matrix population model for brown garden snail (*Cornu aspersa*). The original matrix population model (MPM) has dimensionality n=6, and the aggregated MPM has dimensionality m=3. Balancing lowers the first-row values (transitions into the youngest age groups) and raises the second and third rows (transitions into older groups), according to their reproductive values.

TABLE S1. Details of the 12 age-based matrix population models (MPMs) selected from the COMADRE Animal Matrix Database (version 4.23.3.1) for comparing standard aggregation with elasticity-consistent aggregation. MPMs were selected over a wide range of dimensionalities ($n$; number of age classes), projection intervals ($\Delta t$; time step of model in years), and population growth rates ($\lambda$). Each matrix corresponds to a different taxonomic class. Matrix ID: unique identifier of a MPM in the COMADRE database; Common name: the common name of the associated population; Latin name: species name using binomial nomenclature; Class: taxonomic class of the species; Matrix type: indicates whether the MPM's matrix is primitive (strictly dominant eigenvalue, which guarantees convergence to the stable age distribution) or imprimitive (no strictly dominant eigenvalue, which fosters persistent oscillations); Reference: original publication of the MPM.

| Matrix ID | Common name | Latin name | Taxonomic Class | $n$ | $\Delta t$ | $\lambda$ | Matrix type | Reference |
|---|---|---|---|---|---|---|---|---|
| 240385 | Rocky Mountain elk | Cervus canadensis nelsoni | Mammalia | 23 | 1.00 | 1.05 | Primitive | Clark (2014) |
| 248165 | Chinook salmon | Oncorhynchus tshawytscha | Actinopterygii | 5 | 1.00 | 0.91 | Primitive | Wilson (2003) |
| 248231 | Axolotl | Ambystoma mexicanum | Amphibia | 4 | 0.07 | 1.00 | Primitive | Zambrano et al. (2007) |
| 248634 | White-tailed eagle | Haliaeetus albicilla | Aves | 37 | 1.00 | 0.95 | Primitive | Krüger et al. (2010) |
| 248846 | Soft-shell clam | Mya arenaria | Bivalvia | 11 | 1.00 | 2.39 | Primitive | Brousseau (1978) |
| 248860 | Water flea | Daphnia magna | Branchiopoda | 7 | 0.01 | 1.56 | Primitive | Duchet et al. (2010) |
| 248877 | Earthworm | Eisenia fetida | Clitellata | 7 | 0.02 | 0.47 | Primitive | Santadino et al. (2014) |
| 248914 | Brown garden snail | Cornu aspersa | Gastropoda | 6 | 1.00 | 1.00 | Primitive | Laskowski & Hopkin (1996) |
| 248989 | Bog fritillary | Proclossiana eunomia | Insecta | 5 | 1.00 | 0.91 | Imprimitive | Radchuk et al. (2013) |
| 249063 | Amphipod | Ampelisca abdita | Malacostraca | 7 | 0.03 | 0.90 | Primitive | Kuhn et al. (2002) |
| 249973 | Polychaete | Nephtys incisa | Polychaeta | 5 | 1.00 | 1.34 | Primitive | Zajac & Whitlatch (1989) |
| 250052 | Freshwater crocodile | Crocodylus johnsoni | Reptilia | 50 | 1.00 | 1.01 | Primitive | Smith & Webb (1985) |

TABLE S2. Summary of the performance of two aggregation methods for estimating generation time across 12 matrix population models. Win rate is the proportion of aggregation levels at which the elasticity-consistent (EC) method was closer to the actual value than the standard method (std). MAE (std) and MAE (EC) denote the median absolute error across aggregation levels for the standard and elasticity-consistent estimates, respectively, providing a measure of typical deviation from the actual value. Directional error records the proportion of estimates that were too high (above the actual value), too low (below the actual value), or exact (equal to the actual value) for each method.

| **Population** | **Win rate** | **MAE (std)** | **MAE (EC)** | **Directional error (std)** | | | **Directional error (EC)** | | |
|---|---|---|---|---|---|---|---|---|---|
| | | | | **Too high** | **Too low** | **Exact** | **Too high** | **Too low** | **Exact** |
| Rocky Mountain elk | 0.75 | 0.21 | 0.01 | 0.27 | 0.73 | 0.00 | 0.50 | 0.00 | 0.50 |
| Chinook salmon | 0.88 | 0.17 | 0.00 | 0.25 | 0.75 | 0.00 | 0.25 | 0.00 | 0.75 |
| Axolotl | 0.83 | 0.07 | 0.00 | 0.33 | 0.67 | 0.00 | 0.33 | 0.00 | 0.67 |
| White-tailed eagle | 0.85 | 0.07 | 0.00 | 0.19 | 0.81 | 0.00 | 0.19 | 0.00 | 0.81 |
| Soft-shell clam | 0.85 | 0.30 | 0.04 | 0.40 | 0.60 | 0.00 | 0.50 | 0.00 | 0.50 |
| Water flea | 0.92 | 0.00 | 0.00 | 0.17 | 0.83 | 0.00 | 0.17 | 0.00 | 0.83 |
| Earthworm | 0.75 | 0.00 | 0.00 | 0.83 | 0.17 | 0.00 | 0.33 | 0.00 | 0.67 |
| Brown garden snail | 0.90 | 0.27 | 0.00 | 0.20 | 0.80 | 0.00 | 0.20 | 0.00 | 0.80 |
| Bog fritillary | 0.50 | 0.00 | 0.00 | 0.00 | 0.00 | 1.00 | 0.00 | 0.00 | 1.00 |
| Amphipod | 0.92 | 0.00 | 0.00 | 0.17 | 0.83 | 0.00 | 0.17 | 0.00 | 0.83 |
| Polychaete | 0.88 | 0.16 | 0.06 | 0.25 | 0.75 | 0.00 | 0.50 | 0.00 | 0.50 |
| Freshwater crocodile | 0.95 | 0.04 | 0.00 | 0.06 | 0.94 | 0.00 | 0.10 | 0.00 | 0.90 |
| Total | 0.86 | n.a. | n.a. | 0.20 | 0.77 | 0.03 | 0.24 | 0.00 | 0.76 |

*Note*: The values in the 'Total' row represent the pooled proportions across all populations and aggregation levels. MAE values are n.a. (not applicable) because these errors are not pooled across populations.

TABLE S3. Summary of the performance of two aggregation methods for estimating Demetrius' entropy across 12 matrix population models. Win rate is the proportion of aggregation levels at which the elasticity-consistent (EC) method was closer to the actual value than the standard method (std). MAE (std) and MAE (EC) denote the median absolute error across aggregation levels for the standard and elasticity-consistent estimates, respectively, providing a measure of typical deviation from the actual value. Directional error records the proportion of estimates that were too high (above the actual value), too low (below the actual value), or exact (equal to the actual value) for each method.

| **Population** | **Win rate** | **MAE (std)** | **MAE (EC)** | **Directional error (std)** | | | **Directional error (EC)** | | |
|---|---|---|---|---|---|---|---|---|---|
| | | | | **Too high** | **Too low** | **Exact** | **Too high** | **Too low** | **Exact** |
| Rocky Mountain elk | 0.80 | 0.65 | 0.65 | 0.00 | 1.00 | 0.00 | 0.00 | 1.00 | 0.00 |
| Chinook salmon | 0.13 | 0.05 | 0.21 | 0.00 | 1.00 | 0.00 | 0.00 | 1.00 | 0.00 |
| Axolotl | 0.17 | 0.00 | 0.37 | 0.67 | 0.33 | 0.00 | 0.67 | 0.33 | 0.00 |
| White-tailed eagle | 0.68 | 0.63 | 0.63 | 0.00 | 1.00 | 0.00 | 0.00 | 1.00 | 0.00 |
| Soft-shell clam | 0.75 | 0.58 | 0.44 | 0.10 | 0.90 | 0.00 | 0.10 | 0.90 | 0.00 |
| Water flea | 0.58 | 0.27 | 0.25 | 0.17 | 0.83 | 0.00 | 0.17 | 0.83 | 0.00 |
| Earthworm | 0.42 | 0.29 | 0.31 | 0.67 | 0.33 | 0.00 | 0.67 | 0.33 | 0.00 |
| Brown garden snail | 0.50 | 0.26 | 0.30 | 0.20 | 0.80 | 0.00 | 0.40 | 0.60 | 0.00 |
| Bog fritillary | 0.50 | 0.00 | 0.00 | 0.00 | 0.00 | 1.00 | 0.00 | 0.00 | 1.00 |
| Amphipod | 0.08 | 0.16 | 0.16 | 0.00 | 1.00 | 0.00 | 0.00 | 1.00 | 0.00 |
| Polychaete | 0.63 | 0.37 | 0.29 | 0.25 | 0.75 | 0.00 | 0.25 | 0.75 | 0.00 |
| Freshwater crocodile | 0.60 | 0.67 | 0.67 | 0.00 | 1.00 | 0.00 | 0.00 | 1.00 | 0.00 |
| Total | 0.60 | n.a. | n.a. | 0.06 | 0.91 | 0.03 | 0.07 | 0.90 | 0.03 |

*Note*: The values in the 'Total' row represent the pooled proportions across all populations and aggregation levels. MAE values are n.a. (not applicable) because these errors are not pooled across populations.

TABLE S4. Summary of the performance of two aggregation methods for estimating net reproductive rate across 12 matrix population models. Win rate is the proportion of aggregation levels at which the elasticity-consistent (EC) method was closer to the actual value than the standard method (std). MAE (std) and MAE (EC) denote the median absolute error across aggregation levels for the standard and elasticity-consistent estimates, respectively, providing a measure of typical deviation from the actual value. Directional error records the proportion of estimates that were too high (above the actual value), too low (below the actual value), or exact (equal to the actual value) for each method.

| **Population** | **Win rate** | **MAE (std)** | **MAE (EC)** | **Directional error (std)** | | | **Directional error (EC)** | | |
|---|---|---|---|---|---|---|---|---|---|
| | | | | **Too high** | **Too low** | **Exact** | **Too high** | **Too low** | **Exact** |
| Rocky Mountain elk | 0.75 | 0.01 | 0.00 | 0.27 | 0.73 | 0.00 | 1.00 | 0.00 | 0.00 |
| Chinook salmon | 0.88 | 0.01 | 0.00 | 0.75 | 0.25 | 0.00 | 0.75 | 0.25 | 0.00 |
| Axolotl | 0.83 | 0.01 | 0.00 | 0.67 | 0.33 | 0.00 | 0.67 | 0.33 | 0.00 |
| White-tailed eagle | 0.82 | 0.00 | 0.00 | 0.86 | 0.14 | 0.00 | 0.83 | 0.17 | 0.00 |
| Soft-shell clam | 0.25 | 10.98 | 20.47 | 0.40 | 0.60 | 0.00 | 1.00 | 0.00 | 0.00 |
| Water flea | 0.08 | 0.03 | 0.85 | 0.17 | 0.83 | 0.00 | 1.00 | 0.00 | 0.00 |
| Earthworm | 0.25 | 0.02 | 0.02 | 0.67 | 0.33 | 0.00 | 0.67 | 0.33 | 0.00 |
| Brown garden snail | 0.90 | 0.00 | 0.00 | 0.20 | 0.80 | 0.00 | 0.80 | 0.00 | 0.20 |
| Bog fritillary | 0.50 | 0.00 | 0.00 | 0.00 | 0.00 | 1.00 | 0.00 | 0.00 | 1.00 |
| Amphipod | 0.92 | 0.00 | 0.00 | 0.83 | 0.17 | 0.00 | 0.83 | 0.17 | 0.00 |
| Polychaete | 0.63 | 0.07 | 0.13 | 0.25 | 0.75 | 0.00 | 1.00 | 0.00 | 0.00 |
| Freshwater crocodile | 0.95 | 0.00 | 0.00 | 0.06 | 0.94 | 0.00 | 1.00 | 0.00 | 0.00 |
| Total | 0.76 | n.a. | n.a. | 0.39 | 0.58 | 0.03 | 0.90 | 0.07 | 0.03 |

*Note*: The values in the 'Total' row represent the pooled proportions across all populations and aggregation levels. MAE values are n.a. (not applicable) because these errors are not pooled across populations.

**References (figures and tables)**


Brousseau, D. J. (1978). Population dynamics of the soft-shell clam *Mya arenaria*. *Marine Biology*, 50, 63–71. https://doi.org/10.1007/BF00390542

Clark, D. A. (2014). *Implications of cougar prey selection and demography on population dynamics of elk in northeast Oregon.* PhD dissertation, Oregon State University.

Duchet, C., Coutellec, M. A., Franquet, E., Lagneau, C., & Lagadic, L. (2010). Population-level effects of spinosad and Bacillus thuringiensis israelensis in Daphnia pulex and Daphnia magna: comparison of laboratory and field microcosm exposure conditions. *Ecotoxicology*, 19(7), 1224–1237. https://doi.org/10.1007/s10646-010-0507-y

Krüger, O., Grünkorn, T., & Struwe-Juhl, B. (2010). The return of the white-tailed eagle (*Haliaeetus albicilla*) to northern Germany: modelling the past to predict the future. *Biological Conservation*, 143(3), 710–721. https://doi.org/10.1016/j.biocon.2009.12.010

Kuhn, A., Munns, W. R., Serbst, J., Edwards, P., Cantwell, M. G., Gleason, T., Pelletier, M. C., & Berry, W. (2002). Evaluating the ecological significance of laboratory response data to predict population-level effects for the estuarine amphipod *Ampelisca abdita*. *Environmental Toxicology and Chemistry: An International Journal*, 21(4), 865–874. https://doi.org/10.1002/etc.5620210425

Laskowski, R., & Hopkin, S. P. (1996). Effect of Zn, Cu, Pb, and Cd on fitness in snails (*Helix aspersa*). *Ecotoxicology and Environmental Safety*, 34(1), 59–69. https://doi.org/10.1006/eesa.1996.0045

Radchuk, V., Turlure, C., & Schtickzelle, N. (2013). Each life stage matters: the importance of assessing the response to climate change over the complete life cycle in butterflies.

*Journal of Animal Ecology*, 82(1), 275–285. https://doi.org/10.1111/j.1365-2656.2012.02029.x

Santadino, M., Coviella, C., & Momo, F. (2014). Glyphosate sublethal effects on the population dynamics of the earthworm *Eisenia fetida* (Savigny, 1826). *Water, Air, & Soil Pollution*, 225, 1–8. https://doi.org/10.1007/s11270-014-2207-3

Smith, A. M. A., & Webb, G. J. W. (1985). *Crocodylus johnstoni* in the McKinlay Area, N.T. VIII. A population simulation model. *Australian Wildlife Research*, 12(3), 541–554. https://doi.org/10.1071/wr9850541

Wilson, P. H. (2003). Using population projection matrices to evaluate recovery strategies for Snake River spring and summer Chinook salmon. *Conservation Biology*, 17(3), 782–794. https://doi.org/10.1046/j.1523-1739.2003.01535.x

Zajac, R. N., & Whitlatch, R. B. (1989). Natural and disturbance-induced demographic variation in an infaunal polychaete, *Nephtys incisa*. *Marine Ecology Progress Series*, 57, 89–102. https://doi.org/10.3354/meps057089

Zambrano, L., Vega, E., Herrera, M., L. G., Prado, E., & Reynoso, V. H. (2007). A population matrix model and population viability analysis to predict the fate of endangered species in highly managed water systems. *Animal Conservation*, 10(3), 297–303. https://doi.org/10.1111/j.1469-1795.2007.00105.
